# Detection of prokaryotic-like ribosome exit tunnels within eukaryotic kingdoms

**DOI:** 10.1101/2025.04.08.647849

**Authors:** Shiqi Yu, Artem Kushner, Simcha Srebnik, Khanh Dao Duc

## Abstract

The ribosome exit tunnel is a critical sub-compartment that actively regulates the folding and dynamics of nascent polypeptide chains during protein translation. In this study, we systematically examined tunnel structures of 725 ribosome models obtained through cryo-EM and X-ray crystallography, to quantify structural variations across different species and biological domains. Hierarchical clustering revealed significant geometric differences between prokaryotic and eukaryotic ribosomes, with a surprising discovery: six eukaryotic protist species display tunnel structures remarkably similar to those of archaea and bacteria. By analyzing the sequences and structures of ribosomal components forming the tunnel walls, we identified four specific sequence modifications in ribosomal proteins and ribosomal RNAs (rRNA) responsible for these unique geometric variations, and detected these modifications in additional protist species lacking existing 3D structural data. Overall, our findings highlights some complex evolutionary mechanisms governing ribosomal protein and large subunit rRNA, providing novel insights into the tunnel’s regulatory role in protein translation.

## Introduction

The ribosome is a large biomolecular complex that mediates the translation of proteins from messenger RNA (mRNA). During this process, the growing polypeptide chain passes through a specialized sub-compartment —the exit tunnel— located within the large ribosomal subunit (LSU) that serves as a corridor for the nascent protein, before emerging into the cellular environment. Historically viewed as a passive conduit, accumulating evidence suggests that the tunnel can actively regulate translation dynamics through intermolecular interaction between the ribosomal components at the exit tunnel and the nascent chain [1–5]. Recent advances in cryogenic electron microscopy (cryo-EM) lead to a surge in high-resolution 3D structures of the ribosomes [6], allowing for comparative analyses that have revealed significant geometric variations in exit tunnels across different species and biological domains [7]. Notably, prokaryotic and eukaryotic tunnels exhibit distinct size and shape characteristics that can substantially influence nascent protein chain dynamics and folding mechanisms [7–9].

Intriguingly, some parasite eukaryotic ribosomes were recently shown to present a unique combination of bacterial and eukaryotic features [10–14]. These ribosomes, characterized by shortened ribosomal RNA fragments and modified protein compositions, approach prokaryotic ribosomes in size, while incorporating specialized modifications potentially related to ribosome quality control [14] and protein folding processes [11]. Despite these findings, we still lack a comprehensive understanding of how these structural variations affect the ribosomal exit tunnel.

This question motivated us to extend our previous investigation of the tunnel heterogeneity [7], that was originally limited to 20 manually curated structures. By integrating comprehensive structural data from 725 ribosomal models spanning 34 species, alongside detailed sequence analyses of ribosomal RNA and proteins, we comprehensively mapped and characterized unique exit tunnel modifications across the eukaryotic domain. Upon automating the computational extraction of tunnel geometric features, we found six eukaryotic protist species with archaeal and bacterial-like tunnel structures. Examining the species led us to identify four specific types of modifications of ribosomal proteins and rRNAs, that differently affect the tunnel geometry in all of its subregions. By retrieving and analyzing homologous nucleotides and protein residues across available sequences, we expanded our initial findings to uncover additional protist species within each identified modification category. Finally, we discuss the potential implications of the modifications that we discovered on protein translation dynamics and evolution of the ribosome, and their potential for the development therapeutic strategies for parasitic diseases.

## Materials and Methods

### Ribosome structures and sequences of ribosomal protein and rRNA

We retrieved 725 cytosolic ribosome structures with structural resolution higher than 6 Å from 34 species across all three domains of life to perform tunnel geometry analysis, excluding one archaeal structure (4V6U) with a lower resolution of 6.6 Å. For the sequence analysis of ribosomal proteins and LSU rRNAs, we retrieved 56,416 uL4 sequences; 67,147 uL22 sequences; 66,538 uL23 sequences; 5,755 eL39 sequences; and 46,850 25S-28S rRNA sequences. Each dataset is described in detail below.

#### Ribosome structures

Cryo-EM reconstructions and X-ray crystallography structures of ribosomes were retrieved from the [Protein Data Bank](https://www.rcsb.org/). The corresponding coordinates of the polypeptide transferase center (PTC) were obtained from [RiboXYZ](https://ribosome.xyz/)‘s archive [15].

#### Ribosomal protein sequences

The sequences of ribosomal proteins were obtained from the InterPro database as a `fasta` archive. Uncharacterized (per InterPro annotations) proteins were left out. With the exception of species that exclusively comprise a single sequence, the sequences were modified through the reduction of strains and sequences within the same species to yield a representative consensus sequence. In cases where a species had fewer than three strains, the longest sequence was selected. Conversely, a dumb consensus sequence was generated with a threshold of 0.75 using Biopython package in Python. An ambiguous residue “X” was added when the threshold was not reached.

#### LSU rRNA sequences

The LSU rRNA sequences for all available eukaryotic species were obtained from the [RNAcentral](https://rnacentral.org/) database and filtered as follows: sequences shorter than 2000 nucleotides in length were left out, sequences with RNAcentral warnings (*e*.*g*., *contaminated*) were left out, *“Partial”* sequences (per author annotation) were left out, sequences representative of the same species (per NCBI) were reduced to a single sequence.

### Automated extraction of ribosome exit tunnel geometry

To extract representations of the exit tunnel from the available structures, we refined the protocol proposed by Dao Duc *et al*. [7] and based on MOLE [16] cavity extraction algorithm. For a single ribosome structure, the process consists of finding three landmark sites to guide the algorithm probe, executing the MOLE cavity extraction algorithm and refining the results of the algorithm. Further details are provided in the Supplementary Information (SI).

The tunnel search algorithm was implemented using MOLE2.5 [17]. The tunnel structure was characterized by its 3D centerline coordinates and the associated radius, extracted with Python. Downstream analysis for computing geometric features of the tunnel was done using Pymol and Python custom scripts.

### Pairwise distance metric for tunnel geometrical comparison

To compare the tunnel structures and quantitatively assess the geometric deviations between tunnels across species, we computed the following pairwise distance for each pair of tunnels [7]:

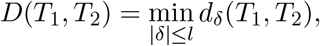

where *T*_1_ and *T*_2_ are two tunnels parametrized by 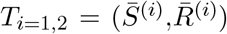 where 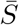 is the arc length parametrization of the tunnel centerline and 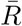 is the associated radius in Å, and

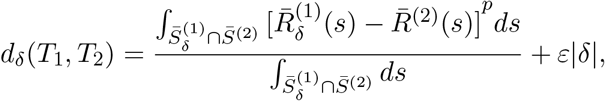

where 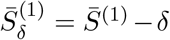 and 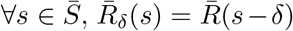. *l* = 20 Å is the maximum shift length, *ε* = 0.01 and *p* = 2 [7]. The minimum distance *D* is a measure of the similarity between two tunnel structures and the *D* values computed for each pair of tunnels comprise the pairwise distance matrix.

### Multiple dimensional scaling and hierarchical clustering of radius plot

To visualize the tunnel structures across species in a lower dimensional space, we ran refined Multidimensional Scaling (MDS) on the pairwise distance matrix, that allowed to jointly detect outliers for a more robust visualization [18]. Subsequently, we refined our analysis by selecting a representative tunnel for each species. This was achieved by identifying the data point closest to the centroid within each species cluster. The associated similarity values between those representative tunnels were then used to generate the hierarchical tree with the Unweighted Pair Group Method Average (UPGMA) algorithm.

### Sequence alignments of ribosomal protein and LSU rRNA

The structure-based sequence alignments of ribosomal proteins and rRNAs were computed by built-in functions `MatchMaker` and `MatchAlign` in Chimera software [19] after superimposition of the molecular structures. The residue-residue distance cutoff was set at 3.5 Å in `MatchAlign`.

The multiple sequence alignments (MSA) of ribosomal proteins were performed by MAFFT [20] with default parameters. The protein sequences were truncated to residues located within 15 Å of the tunnel centerline. For 25S-28S rRNA, the nucleotide sequences were aligned by SINA aligner [21] with the reference LSU datasets from SILVA database. All alignments were visualized and edited in Jalview [22].

### Sequence similarity assessment

To compare the similarity between the truncated uL4 ribosomal protein sequences, we adapted the method proposed by Mervat *et al*. [23], but also accounted for the ambiguous residue “X” in consensus sequences and gaps introduced by MSA. More details are provided in SI.

### Phylogenetic tree generation

To visualize the relationships among the species and their lineages, we used ETE toolkit in Python to construct the tree of eukaryotes based on taxonomy IDs in the NCBI database [24, 25]. The classifications of species of interest are summarized in SI.

To detect the eukaryotic species with similar LSU rRNA sequences, we constructed the phylogenetic tree from sequence alignment of 25S-28S rRNAs by FastTree using the maximum likelihood method [26]. The aligned sequences were truncated to 30nt in length using *S*.*lophii* and *E*.*cuniculi* as references. The nucleotide evolution was analyzed by the generalized time-reversible (GTR) model combined with the “CAT” approximation with 20 rate categories [26]. The tree structures were saved in Newick format and edited in iTOL [27].

## Data availability

The data underlying this article is available in figshare, at https://doi.org/10.6084/m9.figshare.c.7758776.v1.

## Results

### Exit tunnel structure extraction and representation across species

We retrieved 725 high-resolution ribosome structures including 511 from prokaryotes (509 bacteria and 2 archaea) and 214 from eukaryotes, encompassing 14 and 20 species, respectively. Cryo-EM structures comprised 58% of the dataset with an average resolution of 3.35 Å, while the remaining X-ray structures had an average resolution of 3.14 Å. To extract the exit tunnels, we implemented a tunnel search algorithm on the LSU rRNA structure, as depicted in Figure 1a and detailed in the Methods section. The geometric properties of the tunnel were simplified into a coordinate set that describes both the centerline trajectory and the tunnel radius at each centerline position, as illustrated in Figure 1b.

**Figure 1:**
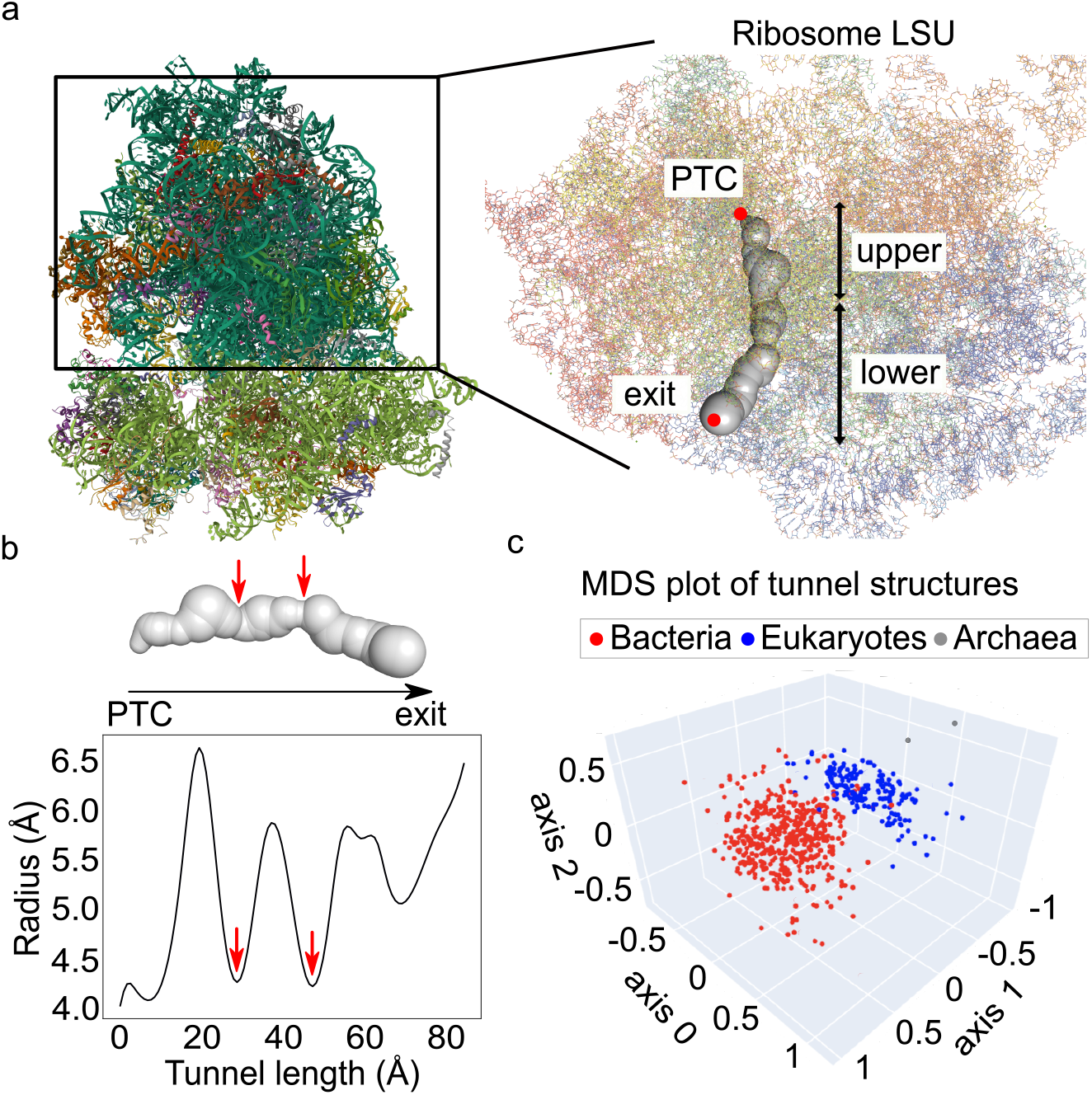
Extraction of the ribosome exit tunnel coordinates and pairwise geometric comparison using radial plots. **(a)** Exit tunnel construction from the large subunit (LSU) rRNA of *H*.*sapiens*. Our developed tunnel searching pipeline locates the peptidyl-transferase center (PTC) and the exit port. The upper region (from the PTC to the constriction site) and the lower region are shown. **(b)** The tunnel structure is stored as radial variation along the centerline, where the PTC is located at *x* = 0 and the constriction sites are marked by red arrows. **(c)** A pairwise distance metric of 725 tunnel structures is transformed into a three-dimensional map using MDS, where each point represents a tunnel structure, with Euclidean distances between points on the map indicating their similarities.

To compare tunnels, we computed distances between radial plots [7] (see Methods), and ran a multi-dimensional scaling (MDS) analysis of the pairwise distance matrix [18], shown in Figure 1c. We observe a clear separation between prokaryotic and eukaryotic tunnels, as previously discovered [7]. To investigate geometric variations at the species level, we selected a representative tunnel for each species, as many species have multiple PDB entries (*e*.*g*., *H*.*sapiens* has 28 strains by cryo-EM). The representative tunnel was defined as the one closest to the centroid of each species’ cluster (detailed examples are provided in Figure S1). The list of 34 selected ribosome structures, including PDB identifiers and resolution details is provided in Table 1.

**Table 1:**
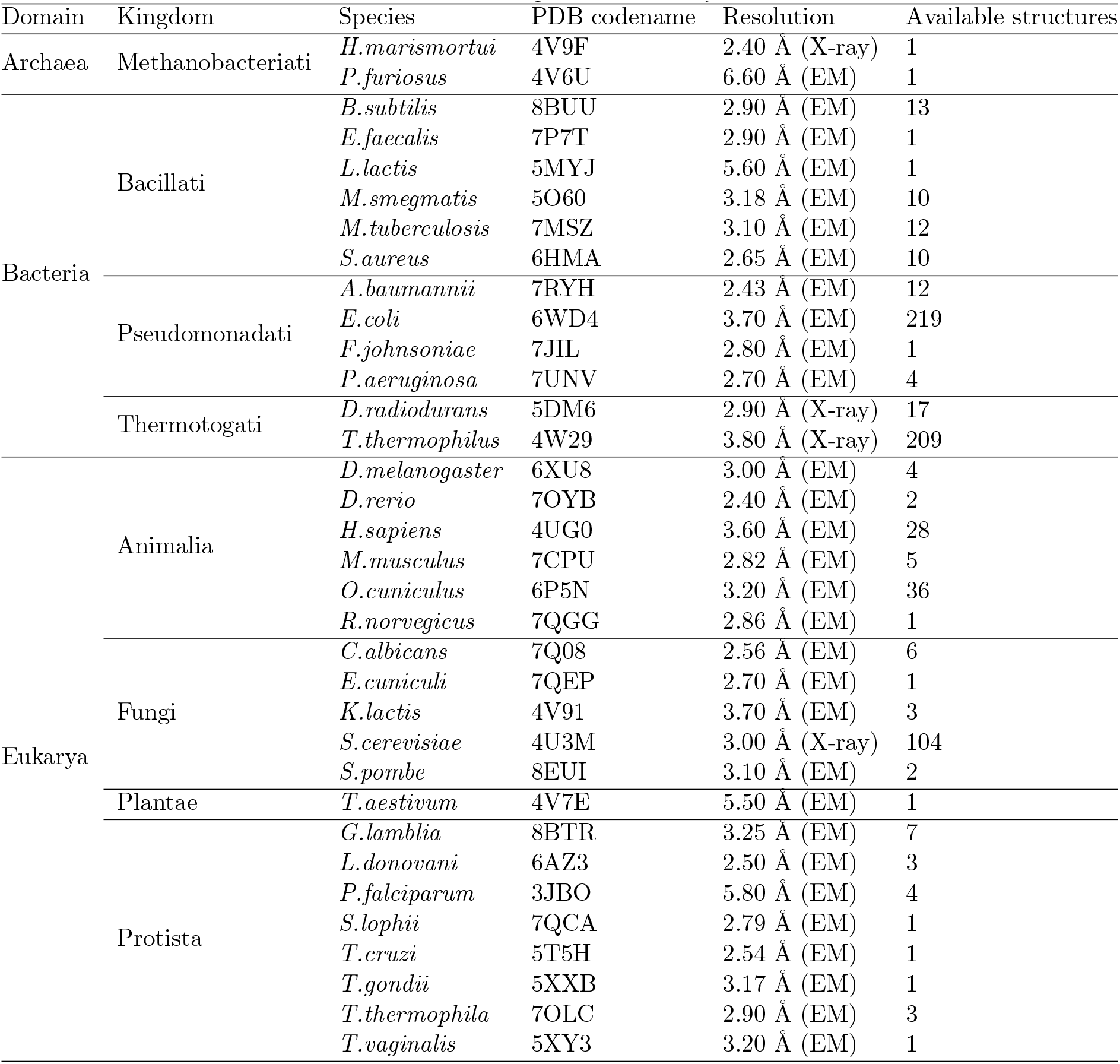
Ribosome structures selected for tunnel geometric analysis with taxonomic classification.

### Hierarchical clustering identifies eukaryotic species with prokaryotic-like tunnel structures

We conducted hierarchical clustering on 34 species based on the geometric similarities of their representative tunnels to visualize the species phylogeny (see Methods). The resulting tree, shown in Figure 2a, is comprised of two primary branches. The upper branch includes most eukaryotes and archaea, while the lower branch predominantly comprises bacterial species. In the upper branch, trypanosomes *T*.*cruzi* and *L*.*donovani* are clustered with two archaeal species, consistent with previous findings [7]. Notably, *T*.*vaginalis* appears separately from other species, and forms an individual branch. In the lower branch, three eukaryotic species, *G*.*lamblia, S*.*lophii* and *E*.*cuniculi*, are grouped with bacteria, where *G*.*lamblia* is embedded in the bacterial cluster, while the other two are grouped separately. Overall, this hierarchical tree highlights six eukaryotic outliers that cluster closer to prokaryotes. To assess the robustness of our results, we compared this clustering with the one generated using the highest-resolution ribosomal structure available for each species (see Figure S2 and Table S1). The consistent clustering patterns suggest that ribosome structures resolved to better than 6 Å reliably preserve tunnel geometric features and do not significantly affect resulting tunnel pairwise comparisons.

**Figure 2:**
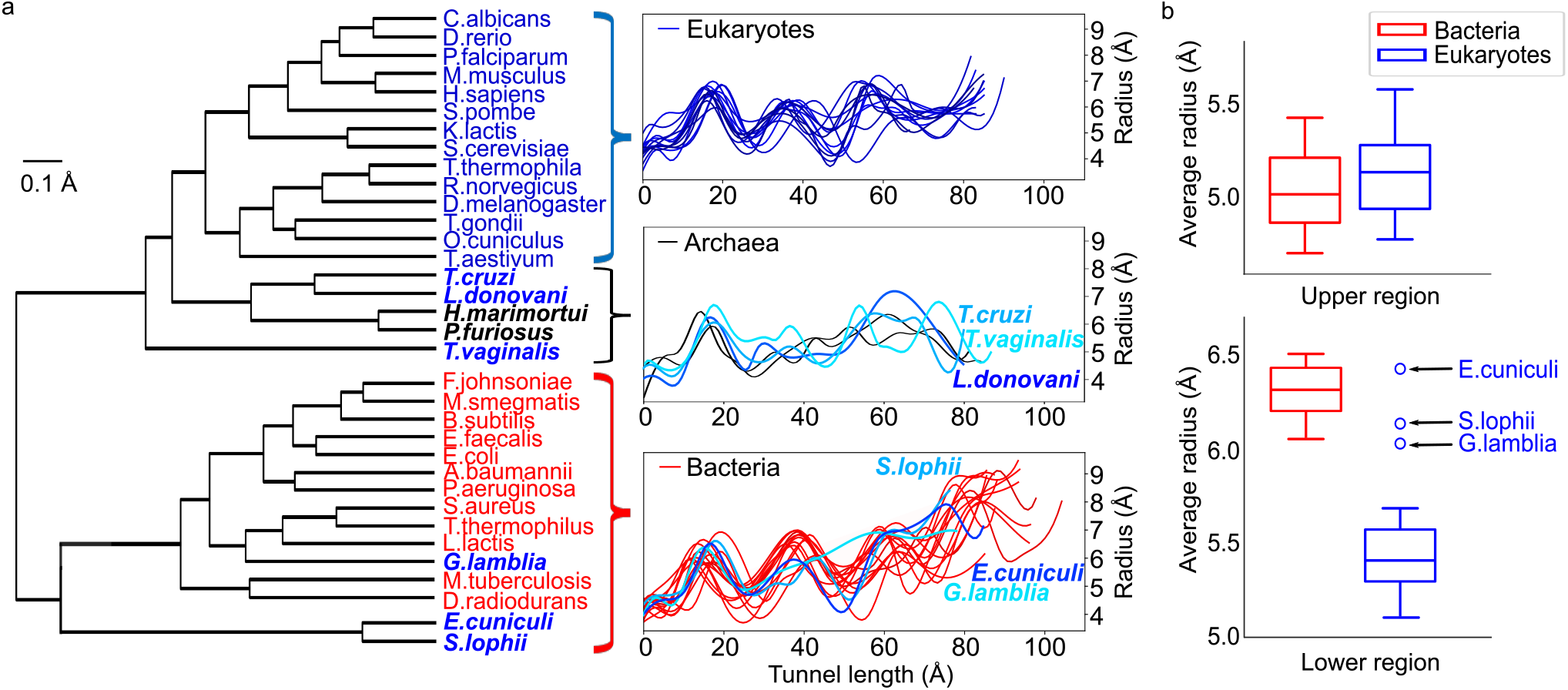
Eukaryotic outliers identified through hierarchical clustering exhibit an average tunnel radius in the lower region comparable to that of bacteria. **(a)** The hierarchical clustering of the representative tunnels constructed based on the pairwise distance matrix. The tree prominently shows six notable eukaryotic species (highlighted in bold): *T*.*cruzi, L*.*donovani, T*.*vaginalis, G*.*lamblia, E*.*cuniculi, S*.*lophii* which are grouped with prokaryotes. Shown on the right are the radial plots corresponding to each species, which are grouped together according to the clustering. **(b)** We compare the average radii of tunnels in the upper and lower regions (up to 80 Å) of species in our dataset. The horizontal line within boxes represents the median value. The outliers observed in the lower region are labeled with their species name.

By examining the radial plots (right panel in Figure 2a), we observe distinct geometric variations in tunnel across species, particularly in the tunnel length and its radius near the exit port. Bacterial tunnels show longer tunnel lengths on average, followed by archaeal and eukaryotic tunnels (see also Figure S3a), in concordance with previous findings [7]. Furthermore, bacterial tunnels, along with the three eukaryotic outliers *E*.*cuniculi, S*.*lophii*, and *G*.*lamblia*, display wider exit ports compared to other eukaryotes. Conversely, the radial plots of *T*.*cruzi, L*.*donovani*, and *T*.*vaginalis* aligned well with those of archaea, and are characterized by narrower exit ports and a greater variation in the radius beyond the first constriction site. These results suggest that the upper tunnel region, extending from the PTC to the constriction site, remains highly conserved across all species. In contrast, the structural divergence originates primarily in the lower tunnel region, from the constriction site to the exit port.

We compared bacterial and eukaryotic tunnels by computing their average radius in the upper and lower regions of the tunnel, as shown in Figure 2b. While the average radii of the upper region of bacteria and eukaryotes are similar, with medians of 5 Å and 5.12 Å, respectively, bacteria show a significantly larger median radius in the lower region of 6.27 Å, compared to 5.31 Å in eukaryotes.

Notably, we identified *E*.*cuniculi, S*.*lophii*, and *G*.*lamblia*, with average tunnel radii comparable to those of bacteria. This may explain the impact of lower regions on clustering results, driven by the expansion of the exit ports in *E*.*cuniculi* and *S*.*lophii*, as well as the enlargement observed in *G*.*lamblia* at the second constriction site, measuring 5.72 Å (Figure S3b).

### Overview of eukaryotic species with structural modifications of ribosome proteins and rRNA

To explain the structural differences in the six eukaryotic outliers whose tunnel structure clusters with prokaryotes, we performed comparative analyses of ribosomal proteins and rRNAs at both structural and sequence levels. We uncovered fours types of tunnel structural modifications and categorized the outliers into four groups: Group A includes *G*.*lamblia* from the phylum *Fornicata*, characterized by a truncated uL4 loop structure. Group B encompasses *T*.*vaginalis*, a parasite from the phylum *Parabasalia*, known as causative agents of trichomoniasis [28], which exhibits single residue modifications in both uL4 and eL39. Group C comprises two trypanosome species, *T*.*cruzi* and *L*.*donovani*, both exhibiting multiple LSU rRNA chains. Lastly, Group D includes *S*.*lophii* and *E*.*cuniculi* from the phylum *Microsporidia*, which are identified by a missing rRNA segment near the tunnel exit port. The corresponding structures for each group are depicted in Figure 3a.

**Figure 3:**
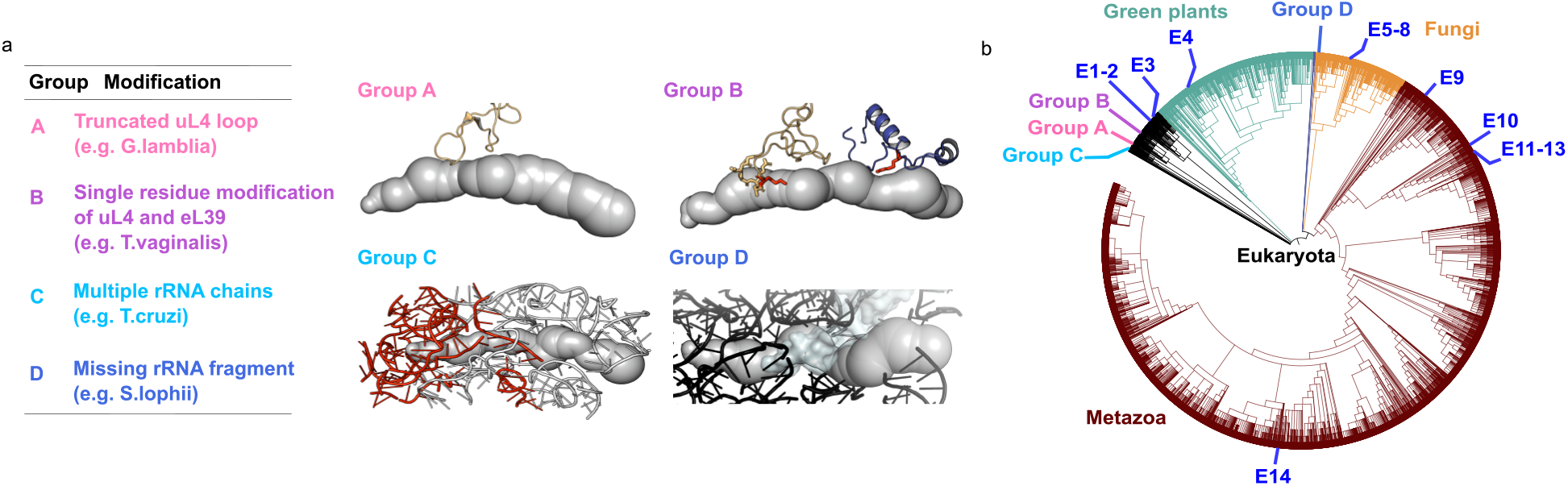
Phylogenetic relationships between eukaryotic species involved in our study with or without tunnel structural modifications. **(a)** Tunnel modifications of eukaryotic outliers are categorized into four groups based on the type of ribosomal protein/RNA modifications. The corresponding structures are depicted on the right. **(b)** The eukaryotic life tree, constructed using taxonomy IDs, highlighting three major clades: green plants (green), fungi (gold), and metazoa (brown). Other smaller clades are shown in black. The evolutionary position of each group of species is labeled on the tree. No tunnel modification is found for species in group E, and the numbering order is consistent with Table S2.

Figure 3b presents the eukaryotic tree of life (eTOL), highlighting the phylogenetic positions of the species studied. Notably, outliers with identical structural modifications are clustered within the same clade, as summarized in Table S2. Furthermore, the sequence analyses of ribosomal proteins and rRNA reveals additional protist species with similar tunnel modifications under each group, as detailed in Table 2. Interestingly, except for the species in Group D, other protists are categorized within the supergroup *Excavata* (see Figure S4) [29]. Their close evolutionary relationships suggests these protists may share commonalities at morphological or physiological levels. We provide detailed analyses of each modification type in the following sections.

**Table 2:**
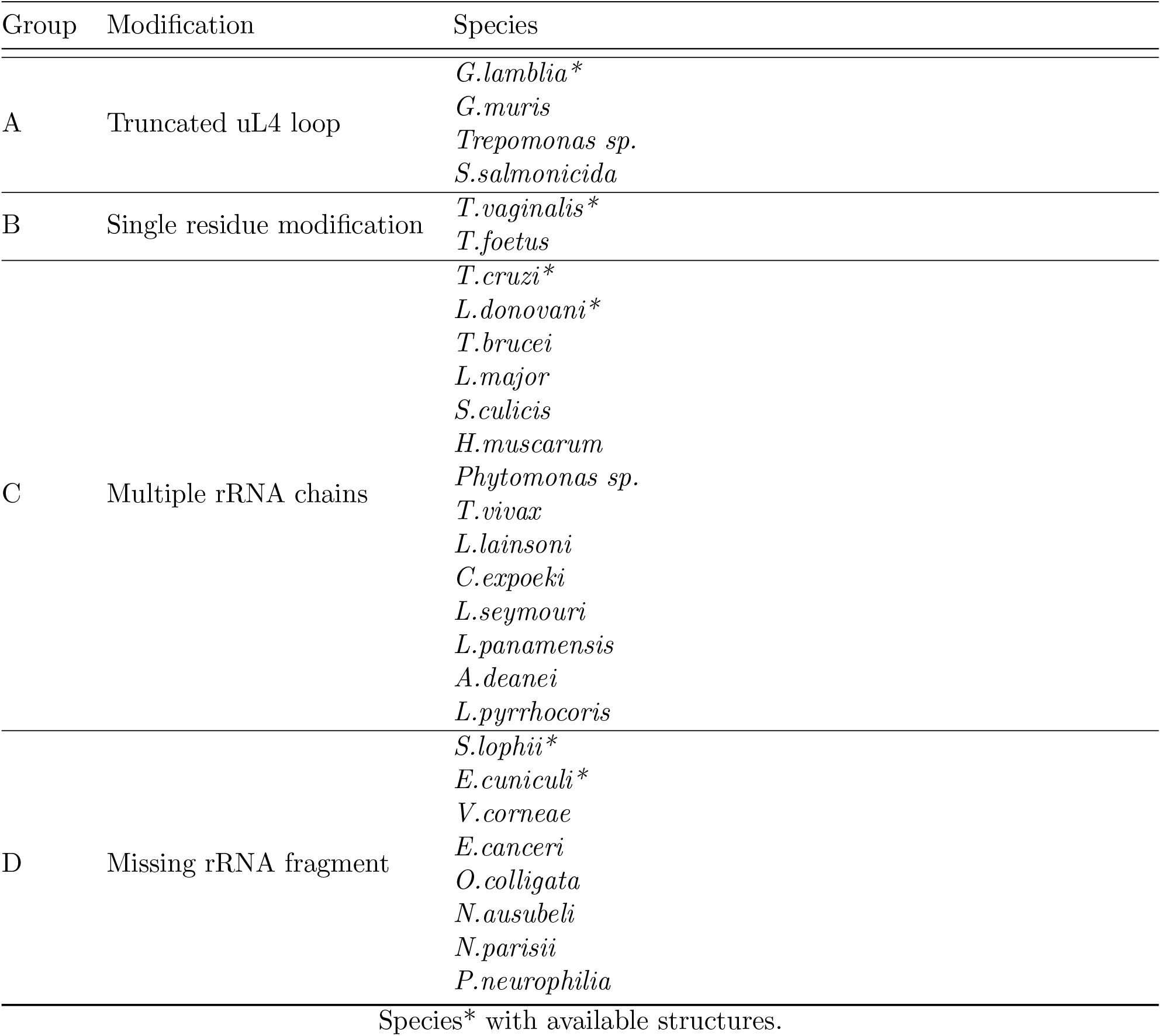
Eukaryotic species identified with ribosomal protein/RNA modifications Group.

### Truncated loop structure of uL4 (Group A)

The tunnel in *G*.*lamblia* exhibits the widest second constriction site among eukaryotes. To explain this anomaly, we superimposed the ribosomal proteins uL4 and uL22, which induce the second constriction site in eukaryotes, and performed structure-based sequence alignments (see Methods section). We found that uL22 is highly conserved close to the tunnel (within 15 Å from centerline) at both structure- and sequence-levels (provided in Figure S5). In contrast, we observed a notable deviation in uL4 sequence of *G*.*lamblia*, with 6 amino acid residues missing, as shown in Figure 4a. The missing residues result in a truncated loop structure of uL4. Note that in bacterial species (*e*.*g*., *E*.*coli*), this loop is completely missing due to the absence of 12 amino acid residues.

**Figure 4:**
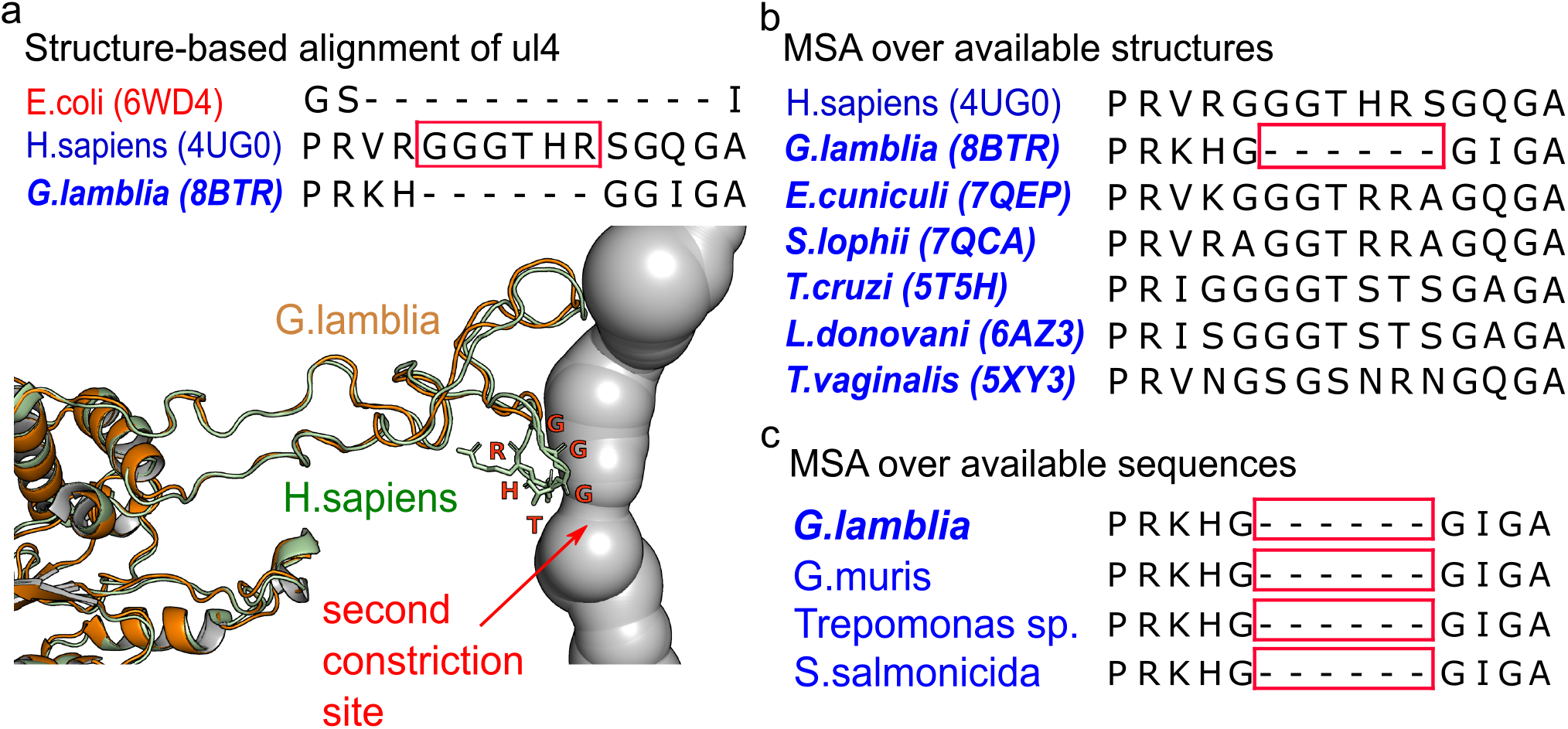
A truncated loop of ul4 protein in *G*.*lamblia* induces a wider second constriction site of the exit tunnel. **(a)** The upper inset shows the structure-based sequence alignment of uL4 proteins for selected species. The gaps represent the structures that are not matched in superposition. The lower inset shows the tunnel structure of *H*.*sapiens* and the structural superposition of uL4 proteins between *G*.*lamblia* (sand) and *H*.*sapiens* (green). The protein structure associated with the missing segment in *G*.*lamblia* are highlighted with the amino acids in red. **(b)** Multiple sequence alignment of uL4 ribosomal proteins for several eukaryotic species. Only the motifs close to the second constriction site of the tunnel (from aligned position 93 to 110) are displayed. **(c)** The alignments of uL4 subsequences from those four species. The important missing segments are highlighted in red boxes.

The divergence of uL4 in *G*.*lamblia* was also observed in the MSA as depicted in Figure 4b.

Notably, only *G*.*lamblia* displays a deletion flanked by two highly conserved glycines. Leveraging these sequence features, we retrieved all the available eukaryotic uL4 sequences to detect species with uL4 structure akin to *G*.*lamblia* but that do not have a ribosome 3D structure. We calculated the dissimilarity scores [23] of all uL4 sequences and compared them to *G*.*lamblia*. We identified three species with a score of zero, indicating similar mutations: *G*.*muris, Trepomonas sp*. and *S*.*salmonicida*. Figure 4c presents their uL4 sequence alignment, highlighting consistent deletion sites without additional mutations. This absence of the conserved “interglycine loop” consequently results in a truncated loop structure of uL4, contributing to a wider second constriction site.

### Specific residue mutations of uL4 and eL39 (Group B)

The radial plots reveal that *T*.*vaginalis* displays a flatter first constriction site and a pronounced third trough at 60 Å from the PTC, which may affect its clustering (see Figure 2). To elucidate the underlying causes of these geometric variations, we focused on the structures of LSU rRNAs and ribosomal proteins (*e*.*g*., uL4, uL22 and eL39) surrounding these regions. The analysis of rRNA revealed no significant impact on tunnel structure, as detailed in Figure S6 and discussed further in the Discussion section. However, MSA of truncated sequences of protein uL4 and eL39 across eukaryotes (Figure 5) highlighted key mutations in *T*.*vaginalis*. Specifically, the highly conserved hydrophobic residues (*e*.*g*., Val (V), Tyr (Y)) at aligned positions 92 in uL4 and 36 in eL39, respectively, are substituted by basic, positively charged residues Lys (K) and Arg (R).

**Figure 5:**
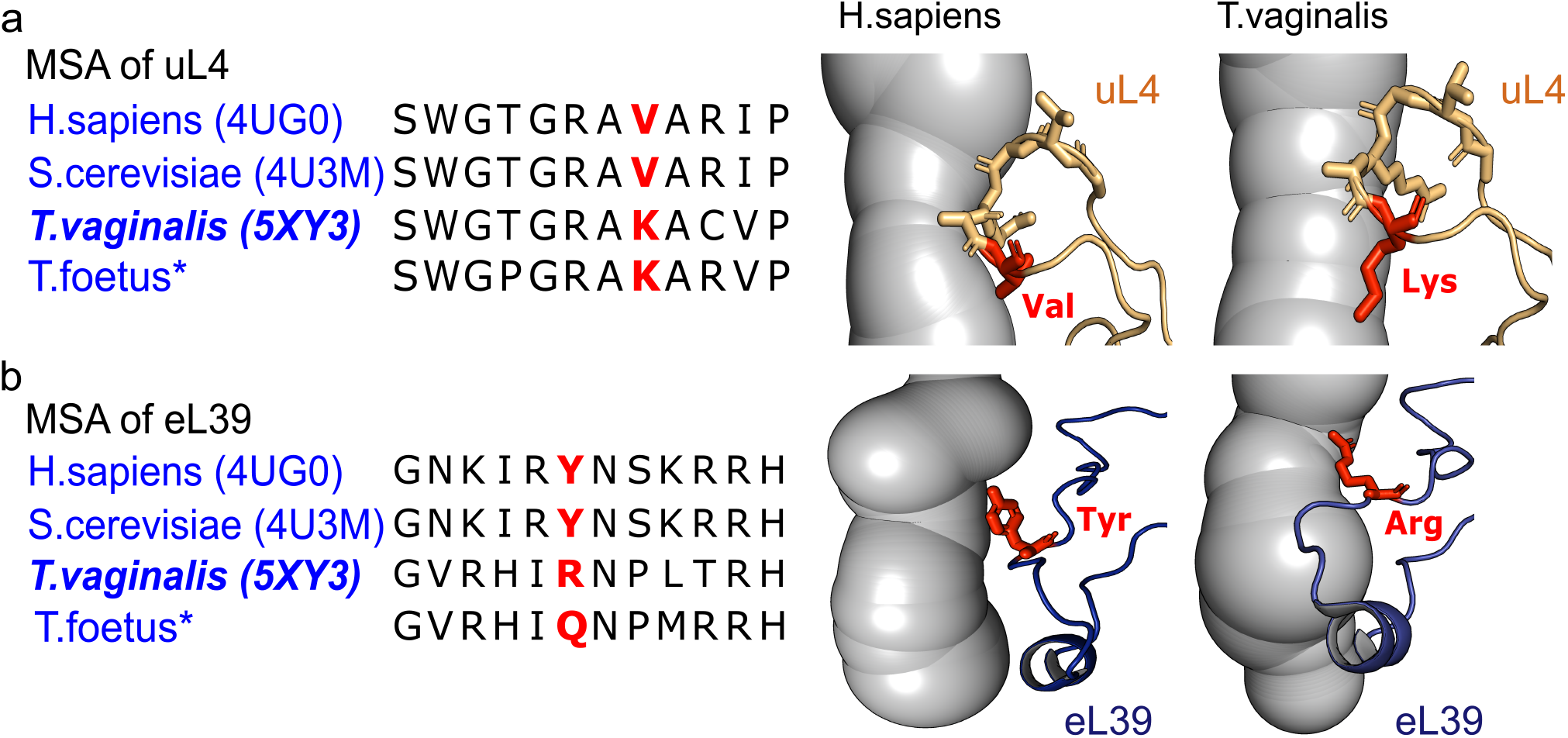
Modifications of single residue in uL4 and eL39 explain the divergent of the tunnel structure in *T*.*vaginalis*. The MSAs of **(a)** uL4 and **(b)** eL39 with the corresponding structure of exit tunnels on the right. The muted residues are marked in red and the longer side chain of non-conserved Lys (K) and Arg (R) alters the tunnel structure in *T*.*vaginalis*. By examining the sequence of uL4 and eL39, *T*.*foetus* is another species that exhibits similar residue mutations at the same sequence sites.

The structural changes associated with these mutated residues are in agreement with the observed differences in tunnel geometry: The longer side chain of Lys aligns parallel to the tunnel surface, generating a looser constraint by uL4, which results in a wider first constriction site of the tunnel. The extended side chain of Arg in eL39, on the other hand, protrudes into the exit tunnel, narrowing down the tunnel compared with other eukaryotes, as shown in Figure 5.

As we did for Group A, we further examined all available sequences of uL4 and eL39, focusing on these specific mutation sites through MSA. Our findings uncovered that only *T*.*foetus* had a Lys mutation in uL4, and none had an Arg mutation in eL39. However, *T*.*foetus* displayed a similar mutation in eL39 at the same aligned position, where the hydrophobic residue was replaced by another basic, positively charged residue Gln (Q). The configuration of Gln side chain suggests that similar to *T*.*vaginalis*, the tunnel may be narrower near the exit port.

### Multiple fragmentation of LSU rRNA (Group C)

The radial plots of *T*.*cruzi* and *L*.*donovani* demonstrated compact constriction sites, which may account for their clustering with archaea (see Figure 2a). To substantiate this observation, we compared the distances between two constriction sites in eukaryotes and archaea, and found that *T*.*cruzi* and *L*.*donovani* exhibit shorter distances, similar to those in archaea (see Figure S7). As sequence analyses of uL4 and uL22 at the constriction site showed no mutations compared to other eukaryotes (see Figure S8 and S5), we further compared the structure of the rRNA. For these two species, the 28S rRNA is divided into two primary fragments: the LSU-*α* and LSU-*β* 251 chains, shown in Figure 6a. The break between these rRNA fragments was thought to be the main factor compressing the tunnel constriction sites. To pinpoint this break position, we identified the nucleotides within 15 Å from the tunnel surface in each chain (see Figure 6b for *T*.*cruzi* and Figure S7b for *L*.*donovani*). The calculation of distances between these nucleotides showed that 31% and 22% of the nucleotide pairs (*≤* 10 Å) converge near the constriction sites in *T*.*cruzi* and *L*.*donovani*, respectively. While the remaining are located around the upper tunnel region, illustrated in Figure6d. The convergence of LSU-*α* and LSU-*β* at these regions supports our hypothesis, highlighting a structural mechanism of rRNA that can reduce the distance between tunnel constriction sites.

**Figure 6:**
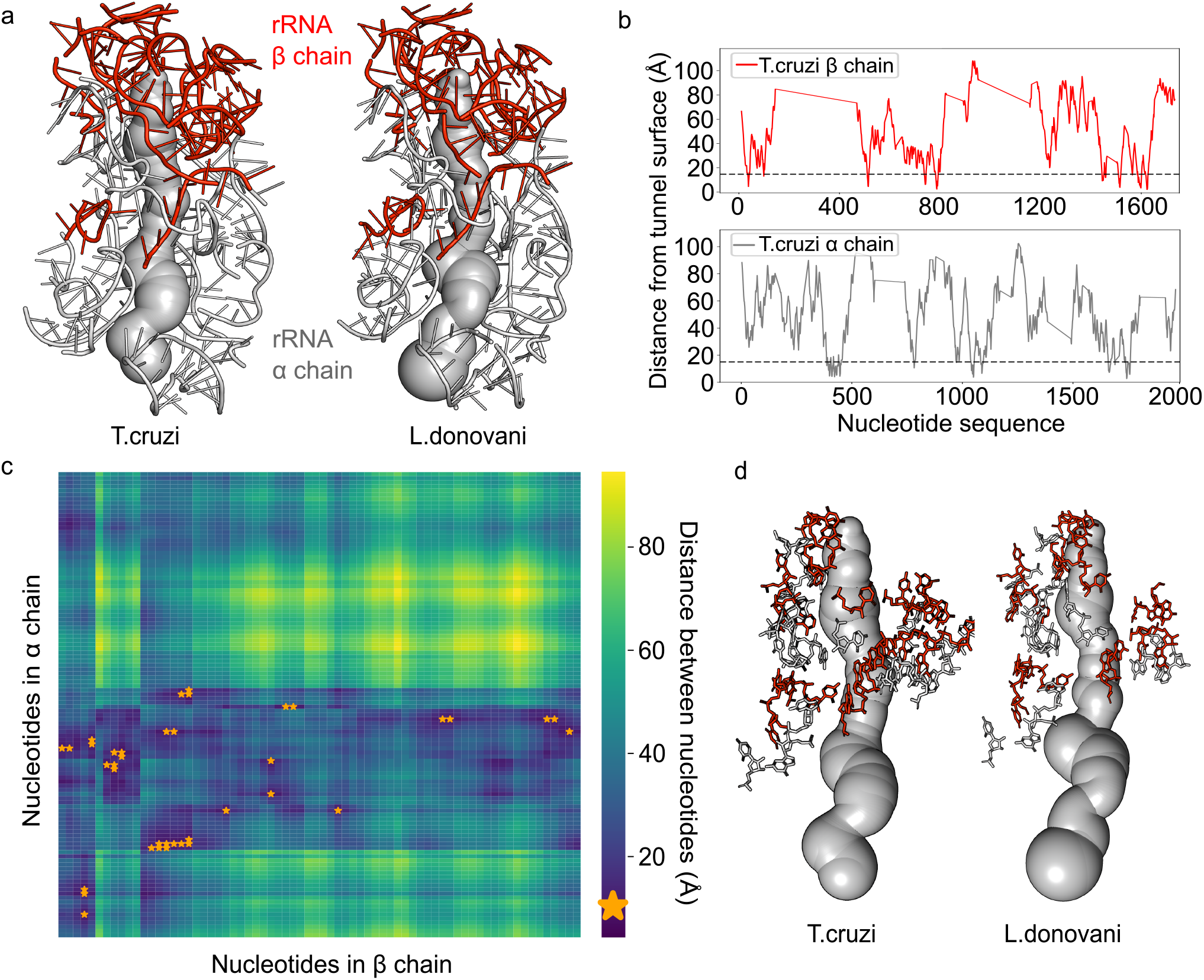
The conjunction of LSU-*α* and LSU-*β* in *T*.*cruzi* and *L*.*donovani* leads to structural variations in constriction sites. **(a)** Tunnel structures with LSU-*α* and LSU-*β* chains of *T*.*cruzi* and *L*.*donovani*. **(b)** We calculate the distances between the tunnel surface and the nucleotides in *β* (upper) and *α* (lower) chains. The spatial position of each nucleotide is calculated at its centroid. Nucleotides within 15 Å of the tunnel surface are selected and the cutoff distance is indicated by the dash lines. **(c)** For *T*.*cruzi*, we map the distances between the selected nucleotides in a heat map. The nucleotide pairs with distance less than 10 Å are highlighted with orange stars. **(d)** The positions of the nucleotides with stars in **(c)**, showing that two rRNA chains meet mostly in the upper region of the tunnel and are close to the constriction sites.

The fragmentation of the LSU rRNA into multiple pieces, referred to as “hidden breaks” [30, 31], has been extensively studied over the decades. Natsidis *et al*. [31] previously developed a computational method to diagnose the presence of the hidden break in 28S rRNA through mapping of RNA-Seq data, and discovered non-homologous hidden breaks in protostomes and vertebrates [31]. Employing this detection pipeline [31–34], we identified breaks in *T*.*cruzi* and *L*.*donovani* at distinct sequence positions than any previously reported (Figure S7d), yet this method did not determine the break location within the LSU. To further identify other organisms with breaks near the tunnel constriction sites, we first conducted a sequence analysis of 25S-28S rRNAs from the eukaryotic species studied, where we observed a conserved motif, 5’-GAAAAGGGGCA-3’ in the *α* chain of both *T*.*cruzi* and *L*.*donovani* which is absent in other species. Its corresponding structure is located at the edge of the break (Figure S7e), suggesting it as a landmark for the *trypanosome*-specific rRNA configuration that could lead to a shorter distance between two constriction sites. Utilizing this sequence signature, we performed the MSA over all available LSU rRNAs and identified 12 other eukaryotic species with this conserved sequence, all belonging to the *Trypanosomatidae* family.

### Missing fragment of rRNA at the tunnel exit port (Group D)

The radial plots of the tunnels suggest that the vestibules of *S*.*lophii* and *E*.*cuniculi*, enclosed by ribosomal protein eL39 and rRNA, are more similar in size to bacteria (see Figure 2). To explain this tunnel modification, we first performed structure-based sequence alignment of eL39 across eukaryotes. The result showed that all sequences were well-aligned and no divergence appeared in *S*.*lophii* and *E*.*cuniculi* (see Figure S9). Consequently, we compared LSU rRNA structures. To reduce the computational costs, we isolated the 25S-28S rRNA structures, restricting our analysis to residues located within a 30 Å radius of the tunnel centerline. Superimposition of these edited rRNA structures revealed that *S*.*lophii* and *E*.*cuniculi* exhibit a deletion of approximately 10 nucleotides compared to other eukaryotes as shown in Figure 7a (complete MSA is provided in Figure S10). This missing segment results in an expanded exit port opening, as depicted in Figure 7b (see *E*.*cuniculi* in Figure S10c), possibly underlying its separation from the upper branch in the clustering.

**Figure 7:**
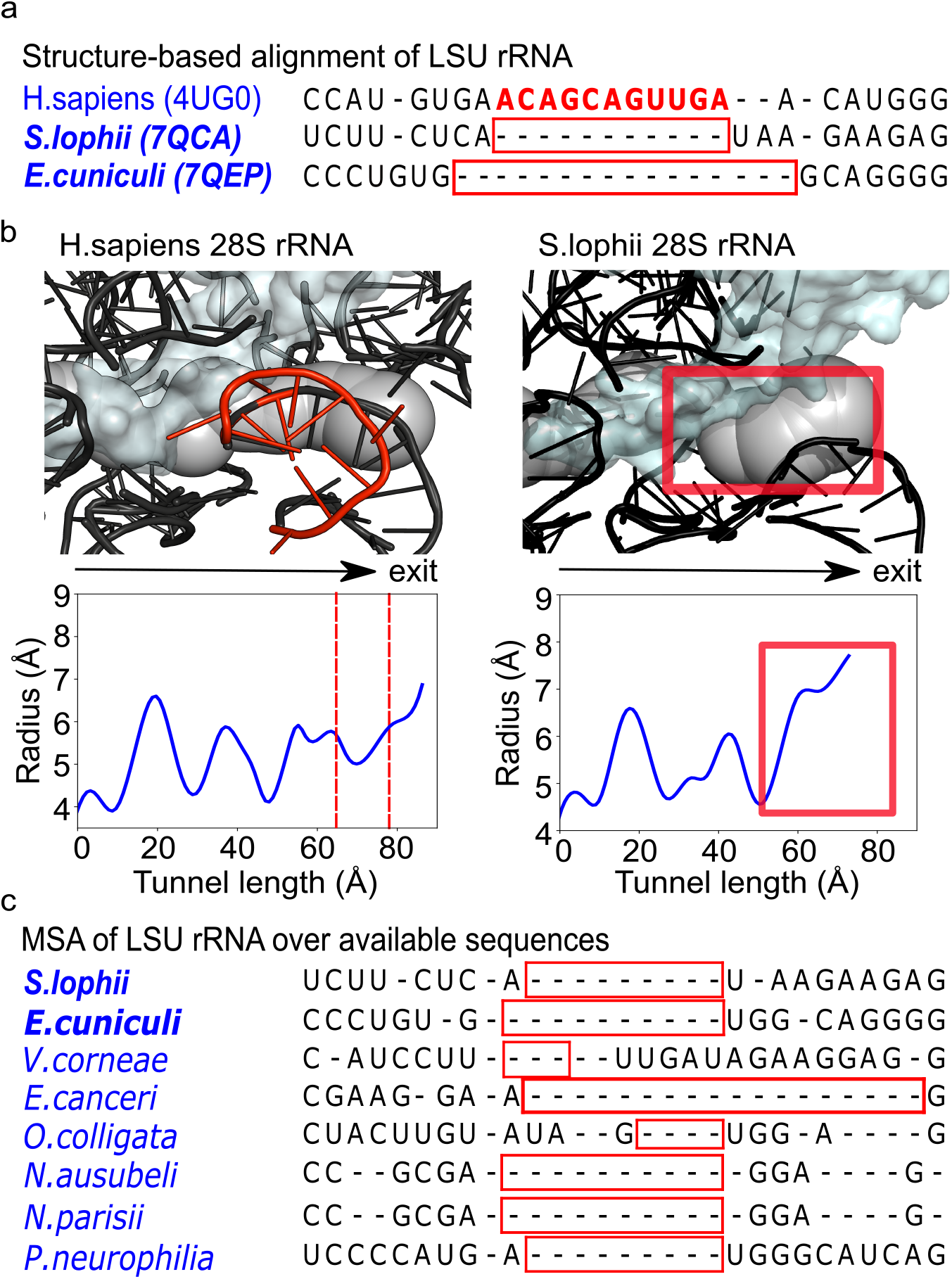
The missing fragment of rRNA at the tunnel exit port explains the larger size of vestibules in *S*.*lophii* and *E*.*cuniculi*. **(a)** Structure-based alignment of LSU rRNA subsequences from eukaryotes. Nucleotides highlighted in red correspond to structures that failed to be matched during superposition. **(b)** The comparisons of LSU rRNA structures and associated radial plots between *H*.*sapiens* and *S*.*lophii*. The red cartoon structure in *H*.*sapiens* relates to the highlighted nucleotides in **(a)**, and its position is enclosed by dash lines in the radial plot. The missing counterpart in *S*.*lophii* is indicated by red box, where a large opening can be observed. **(c)** The MSA of LSU rRNA from the eukaryotic species clustered with *E*.*cuniculi* and *S*.*lophii* in the phylogenetic tree from the alignment of LSU rRNA. Species with missing segments, highlighted in red boxes, exhibit length differences compared to *E*.*cuniculi* and *S*.*lophii*. These variations may result in vestibules of different sizes.

To identify additional eukaryotic species with such modifications, we constructed a phylogenetic tree based on LSU rRNA sequence alignments (see Methods) to group species with conserved sequence features [35]. We found that *S*.*lophii* and *E*.*cuniculi* cluster within the same subtree, alongside six other species, summarized in Table 2. As shown in Figure 7c, *N*.*ausubeli, N*.*parisii*, and *P*.*neurophilia* exhibit higher alignment scores with both *E*.*cuniculi* and *S*.*lophii*, evident in the consistent placement of gaps and nucleotides. In contrast, *V*.*corneae* and *O*.*colligata* show shorter missing segments, while *E*.*canceri* displays longer ones. These sequence variations suggest discrepancies in the tunnel vestibule radius due to varying degrees of deletion of rRNA.

## Discussion

In this study, we investigated the structural heterogeneity of ribosome exit tunnels across various species. Upon comparing tunnel geometries, we identified six eukaryotic protist species that display archaeal and bacterial-like tunnel structures, in contrast to the results of a previous study [7]. This novel finding suggests that protist species have more variability in their ribosome structures, with variations spanning all sub-regions of the ribosome, including the conserved core, constriction site, and exit port. It would be interesting to expand our knowledge of structures of protist ribosomes to potentially uncover more differences, and potentially guide the development of ribosome targeting therapeutic strategies for parasitic diseases [36].

To run our comparative analysis over hundreds of ribosome structures, we automated the extraction of tunnel geometric features by leveraging conserved landmarks, without the need for manual curation. As more high-resolution ribosomes become available, this pipeline can be applied to even larger datasets and facilitate the high-throughput analysis of tunnel structures for more species. Furthermore, our study shows how using sequence alignments can help predict structural features without available structures, and identify protists with closer evolutionary position in eTOL that should display similar tunnel geometries. This highlights the potential of using tunnel structure as a tool for tracing evolutionary lineages and study ribosome evolution.

### Impact of tunnel modifications on translation dynamics in protists

Our study shows that protist species can carry modifications at the constriction site as well as at the exit port of the exit tunnel. As species-specific tunnel modifications at constriction sites may affect polypeptide translation dynamics and ribosome stalling [37], the presence of an additional constriction site in eukaryotes, compared to prokaryotes, can also affect peptide transit and folding [8, 38]. In particular, the wider second constriction site in *giardia*-species is expected to facilitate rapid escape from the tunnel, reducing kinetic trapping of the nascent chain. Moreover, the truncated structure in uL4 may alter the eukaryotic regulatory system that interacts with human cytomegalovirus (hCMV)- and fungal arginine attenuator peptide (AAP)-nascent chains through the uL4 loop, thereby stalling and halting ribosome elongation [1]. The reduced tunnel confinement could also increase their sensitivity to antibiotics (*e*.*g*., metronidazole) or other deleterious compounds, causing irreversible accumulation of deleterious mutations and reduction of parasite genomes [10, 12, 39]. This unique wider region in *G*.*lamblia* appears to be an excellent target site for developing *giardia*-specific drugs to combat giardiasis [13] that minimize side effects to host by enhancing the selectivity and specificity of antiparasitic agents.

On the other hand, tunnel modifications at the exit port may impact peptide folding and chaperone engagement [40]. In the vestibular region, the nascent chain can adopt a *β*-hairpin structure or even a tertiary structure [41] with the help of downstream chaperones to prevent misfolding and aggregation [42]. A narrow exit port in *T*.*vaginalis* may reduce the flexibility and compactness of the nascent chain, and its increased interactions with the polypeptide likely alters the mechanisms of peptide stabilization within the tunnel [43, 44]. In contrast, the enlarged exit port observed in *S*.*lophii* and *E*.*cuniculi* may facilitate the release of nascent chains, potentially permitting an earlier folding onset [45, 46]. Consequently, the altered folding conformations could influence the binding with downstream chaperones and targeting factors [42, 47, 48]. Overall, although the functional implications of each tunnel modification remain to be fully elucidated, the presence of prokaryotic-like tunnel structures may contribute to the adaptation of protists to their unique protein synthesis requirements, environmental conditions and lifestyle [43, 49].

### Three evolutionary scenarii altering the tunnel geometry

Our study suggests the existence of three evolutionary processes of ribosome proteins and rRNAs that alter the structure and geometry of the exit tunnel. First, although the ribosome core structure is universally conserved among all life domains, as evidenced by a high degree of geometric conservation in the tunnel upper region [50, 51], we found that multiple rRNA fragmentations can lead to variations in tunnel constriction sites. Non-homologous breaks in trypanosomes and other eukaryotes indicate divergent evolution of 25S-28S rRNA within the eukaryotic domain. Only the breaks evolved in trypanosomes presumably affects tunnel structure under selective pressure. Secondly, the fact that *giardia*-species display a truncated loop structure in uL4 at the constriction site that was similarly found in archaea, suggests that the uL4 loop structure was acquired progressively during evolution from simpler organisms to more complex eukaryotes. Interestingly, it was also suggested that these variations in conserved ribosomal proteins can reflect the adaptation to specific environments and lifestyles of species [49]. Finally, we can relate our observation of the missing fragment of rRNA at the tunnel exit port to the evolution of rRNA segments. As organisms evolved, the rRNA segments accreted onto the surface of core structure, driving the size variability in LSU rRNAs, with the trend: bacteria *≤* archaea *<* eukaryotes, as proposed by Petrov *et al*. [51]. The intermediate size of protist LSU rRNA between bacteria and eukaryotes suggests an underlying evolutionary pressure for the emergence of rRNA expansions. The lack of specific rRNA fragment in protists may facilitate a more streamlined co-translational chaperone system [11].

### Limitations and future directions

Our comparative analysis of the tunnel geometry relies on analyzing the radial plots. While this representation captures the most important geometric variations of the tunnel [5], it would be interesting to compare the full shapes and use them to produce a phylogenetic tree that would allow us to more specifically infer the evolution of the ribosome exit tunnel [51]. Yet, we note that this approach is usually done at the macroscopic level and involves the choice of morphometrics that can also prove to be challenging [52, 53]. It would also be interesting to directly study the impact of the geometric variations of the tunnel using molecular dynamics (MD) simulations of the nascent chain. While various approaches have been proposed to model the ribosome nascent chain complex [9, 54], modeling the elongation of the nascent chain at the time scale of translation [5] would require some extended coarse grained modeling of the tunnel and ribosome surface, as well as some further processing of the tunnel structures of interest. We are currently pursuing these two directions.

## Author Contributions

SS and KDD designed the research; SY, AK, SS and KDD contributed analytic tools and analyzed data; SY, AK, SS and KDD wrote the article.

## Declaration of Interests

The authors declare no competing interests.

## Acknowledgments

This research is supported by NSERC Discovery Grants RGPIN-2020-05348 and RGPIN-2024-04666. We thank Anton Petrov for his feedback.

## Notes

https://doi.org/10.6084/m9.figshare.c.7758776.v1

### Competing Interest Statement

The authors have declared no competing interest.

### Summary of Updates

We added some discussion on structure resolution and verified the robustness of the hierarchical tree. We also fixed some typos.

## References

[1] Wilson, D. N. and Cate, J. H. D. (2012) The structure and function of the eukaryotic ribosome. Cold Spring Harbor perspectives in biology, 4(5), a011536.

[2] O’Brien, E. P., Hsu, S.-T. D., Christodoulou, J., Vendruscolo, M., and Dobson, C. M. (2010) Transient tertiary structure formation within the ribosome exit port. Journal of the American Chemical Society, 132(47), 16928–16937.

[3] Javed, A., Christodoulou, J., Cabrita, L. D., and Orlova, E. V. (2017) The ribosome and its role in protein folding: looking through a magnifying glass. Acta Crystallographica Section D: Structural Biology, 73(6), 509–521.

[4] Nissley, D. A., Vu, Q. V., Trovato, F., Ahmed, N., Jiang, Y., Li, M. S., and O’Brien, E. P. (2020) Electrostatic interactions govern extreme nascent protein ejection times from ribosomes and can delay ribosome recycling. Journal of the American Chemical Society, 142(13), 6103– 6110.

[5] Dao Duc, K. and Song, Y. S. (2018) The impact of ribosomal interference, codon usage, and exit tunnel interactions on translation elongation rate variation. PLoS genetics, 14(1), e1007166.

[6] Poitevin, F., Kushner, A., Li, X., and Dao Duc, K. (2020) Structural heterogeneities of the ribosome: new frontiers and opportunities for cryo-EM. Molecules, 25(18), 4262.

[7] Dao Duc, K., Batra, S. S., Bhattacharya, N., Cate, J. H., and Song, Y. S. (2019) Differences in the path to exit the ribosome across the three domains of life. Nucleic acids research, 47(8), 4198–4210.

[8] Yu, S., Srebnik, S., and Dao Duc, K. (2023) Geometric differences in the ribosome exit tunnel impact the escape of small nascent proteins. Biophysical Journal, 122(1), 20–29.

[9] Chwastyk, M. and Cieplak, M. (2021) Nascent folding of proteins across the three domains of life. Frontiers in Molecular Biosciences, 8, 692230.

[10] Querido, J. B. (2024) A glimpse into Giardia lamblia unique translational machinery. Structure, 32(4), 377–379.

[11] Li, Z., Guo, Q., Zheng, L., Ji, Y., Xie, Y.-T., Lai, D.-H., Lun, Z.-R., Suo, X., and Gao, N. (2017) Cryo-EM structures of the 80S ribosomes from human parasites Trichomonas vaginalis and Toxoplasma gondii. Cell research, 27(10), 1275–1288.

[12] Nicholson, D., Salamina, M., Panek, J., Helena-Bueno, K., Brown, C. R., Hirt, R. P., Ranson, N. A., and Melnikov, S. V. (2022) Adaptation to genome decay in the structure of the smallest eukaryotic ribosome. Nature communications, 13(1), 591.

[13] Hiregange, D.-G., Rivalta, A., Bose, T., Breiner-Goldstein, E., Samiya, S., Cimicata, G., Kulakova, L., Zimmerman, E., Bashan, A., Herzberg, O., et al. (2022) Cryo-EM structure of the ancient eukaryotic ribosome from the human parasite Giardia lamblia. Nucleic acids research, 50(3), 1770–1782.

[14] Eiler, D. R., Wimberly, B. T., Bilodeau, D. Y., Taliaferro, J. M., Reigan, P., Rissland, O. S., and Kieft, J. S. (2024) The Giardia lamblia ribosome structure reveals divergence in several biological pathways and the mode of emetine function. Structure,.

[15] Kushner, A., Petrov, A. S., and Dao Duc, K. (2023) RiboXYZ: a comprehensive database for visualizing and analyzing ribosome structures. Nucleic Acids Research, 51(D1), D509–D516.

[16] Sehnal, D., Svobodová Vařeková, R., Berka, K., Pravda, L., Navrátilová, V., Banáš, P., Ionescu, C.-M., Otyepka, M., and Koča, J. (2013) MOLE 2.0: advanced approach for analysis of biomacromolecular channels. Journal of cheminformatics, 5, 1–13.

[17] Berka, K., Hanák, O., Sehnal, D., Banáš, P., Navratilova, V., Jaiswal, D., Ionescu, C.-M., Svobodová Vařeková, R., Koča, J., and Otyepka, M. (2012) MOLE online 2.0: Interactive web-based analysis of biomacromolecular channels. Nucleic acids research, 40(W1), W222– W227.

[18] Li, W., Mirone, J., Prasad, A., Miolane, N., Legrand, C., and Dao Duc, K. (2023) Orthogonal outlier detection and dimension estimation for improved MDS embedding of biological datasets. Frontiers in Bioinformatics, 3, 1211819.

[19] Pettersen, E. F., Goddard, T. D., Huang, C. C., Couch, G. S., Greenblatt, D. M., Meng, E. C., and Ferrin, T. E. (2004) UCSF Chimera—a visualization system for exploratory research and analysis. Journal of computational chemistry, 25(13), 1605–1612.

[20] Katoh, K. and Standley, D. M. (2013) MAFFT multiple sequence alignment software version 7: improvements in performance and usability. Molecular biology and evolution, 30(4), 772–780.

[21] Pruesse, E., Peplies, J., and Glöckner, F. O. (2012) SINA: accurate high-throughput multiple sequence alignment of ribosomal RNA genes. Bioinformatics, 28(14), 1823–1829.

[22] Waterhouse, A. M., Procter, J. B., Martin, D. M., Clamp, M., and Barton, G. J. (2009) Jalview Version 2—a multiple sequence alignment editor and analysis workbench. Bioinformatics, 25(9), 1189–1191.

[23] Abo-Elkhier, M. M., Abd Elwahaab, M. A., Abo El Maaty, M.I., et al. (2019) Measuring similarity among protein sequences using a new descriptor. BioMed research international, 2019.

[24] Huerta-Cepas, J., Serra, F., and Bork, P. (2016) ETE 3: reconstruction, analysis, and visualization of phylogenomic data. Molecular biology and evolution, 33(6), 1635–1638.

[25] Schoch, C. L., Ciufo, S., Domrachev, M., Hotton, C. L., Kannan, S., Khovanskaya, R., Leipe, D., Mcveigh, R., O’Neill, K., Robbertse, B., et al. (2020) NCBI Taxonomy: a comprehensive update on curation, resources and tools. Database, 2020, baaa062.

[26] Price, M. N., Dehal, P. S., and Arkin, A. P. (2010) FastTree 2–approximately maximum-likelihood trees for large alignments. PloS one, 5(3), e9490.

[27] Letunic, I. and Bork, P. (2024) Interactive Tree of Life (iTOL) v6: recent updates to the phylogenetic tree display and annotation tool. Nucleic Acids Research, p. gkae268.

[28] Yao, C. and Köster, L. S. (2015) Tritrichomonas foetus infection, a cause of chronic diarrhea in the domestic cat. Veterinary research, 46, 1–16.

[29] Burki, F., Roger, A. J., Brown, M. W., and Simpson, A. G. (2020) The new tree of eukaryotes. Trends in ecology & evolution, 35(1), 43–55.

[30] Applebaum, S. W., Ebstein, R., and Wyatt, G. (1966) Dissociation of ribosomal ribonucleic acid from silkmoth pupae by heat and dimethylsulfoxide: Evidence for specific cleavage points. Journal of Molecular Biology, 21(1), 29–41.

[31] Natsidis, P., Schiffer, P. H., Salvador-Martínez, I., and Telford, M. J. (2019) Computational discovery of hidden breaks in 28S ribosomal RNAs across eukaryotes and consequences for RNA Integrity Numbers. Scientific Reports, 9(1), 19477.

[32] Bray, N. L., Pimentel, H., Melsted, P., and Pachter, L. (2016) Near-optimal probabilistic RNA-seq quantification. Nature biotechnology, 34(5), 525–527.

[33] Li, H., Handsaker, B., Wysoker, A., Fennell, T., Ruan, J., Homer, N., Marth, G., Abecasis, G., Durbin, R., and Subgroup, . G. P. D. P. (2009) The sequence alignment/map format and SAMtools. bioinformatics, 25(16), 2078–2079.

[34] Quinlan, A. R. and Hall, I. M. (2010) BEDTools: a flexible suite of utilities for comparing genomic features. Bioinformatics, 26(6), 841–842.

[35] Gupta, R. S. (1998) Protein phylogenies and signature sequences: a reappraisal of evolutionary relationships among archaebacteria, eubacteria, and eukaryotes. Microbiology and Molecular Biology Reviews, 62(4), 1435–1491.

[36] Jia, X., He, X., Huang, C., Li, J., Dong, Z., and Liu, K. (2024) Protein translation: biological processes and therapeutic strategies for human diseases. Signal Transduction and Targeted Therapy, 9(1), 44.

[37] Wilson, D. N., Arenz, S., and Beckmann, R. (2016) Translation regulation via nascent polypeptide-mediated ribosome stalling. Current Opinion in Structural Biology, 37, 123–133.

[38] Bui, P. T. and Hoang, T. X. (2016) Folding and escape of nascent proteins at ribosomal exit tunnel. The Journal of chemical physics, 144(9).

[39] Majumdar, S., Emmerich, A., Krakovka, S., Mandava, C. S., Svärd, S. G., and Sanyal, S. (2023) Insights into translocation mechanism and ribosome evolution from cryo-EM structures of translocation intermediates of Giardia intestinalis. Nucleic acids research, 51(7), 3436–3451.

[40] Han, Y., Gao, X., Liu, B., Wan, J., Zhang, X., and Qian, S.-B. (2014) Ribosome profiling reveals sequence-independent post-initiation pausing as a signature of translation. Cell research, 24(7), 842–851.

[41] Komar, A. A. (2022) From Alpha to Beta–a co-translational way to fold?. Cell Cycle, 21(16), 1663–1666.

[42] Wilson, D. N. and Beckmann, R. (2011) The ribosomal tunnel as a functional environment for nascent polypeptide folding and translational stalling. Current opinion in structural biology, 21(2), 274–282.

[43] Mishra, R. K., Sharma, P., Khaja, F. T., Uday, A. B., and Hussain, T. (2024) Cryo-EM structure of wheat ribosome reveals unique features of the plant ribosomes. Structure, 32(5), 562–574.

[44] Kišonaitė, M., Wild, K., Lapouge, K., Ruppert, T., and Sinning, I. (2022) High-resolution structures of a thermophilic eukaryotic 80S ribosome reveal atomistic details of translocation. Nature Communications, 13(1), 476.

[45] Ahn, M., Wlodarski, T., Mitropoulou, A., Chan, S. H., Sidhu, H., Plessa, E., Becker, T. A., Budisa, N., Waudby, C. A., Beckmann, R., et al. (2022) Modulating co-translational protein folding by rational design and ribosome engineering. nature communications, 13(1), 4243.

[46] Samatova, E., Komar, A. A., and Rodnina, M. V. (2024) How the ribosome shapes cotranslational protein folding. Current Opinion in Structural Biology, 84, 102740.

[47] Vainberg, I. E., Lewis, S. A., Rommelaere, H., Ampe, C., Vandekerckhove, J., Klein, H. L., and Cowan, N. J. (1998) Prefoldin, a chaperone that delivers unfolded proteins to cytosolic chaperonin. Cell, 93(5), 863–873.

[48] Han, P., Shichino, Y., Schneider-Poetsch, T., Mito, M., Hashimoto, S., Udagawa, T., Kohno, K., Yoshida, M., Mishima, Y., Inada, T., et al. (2020) Genome-wide survey of ribosome collision. Cell Reports, 31(5).

[49] Melnikov, S., Manakongtreecheep, K., and Söll, D. (2018) Revising the structural diversity of ribosomal proteins across the three domains of life. Molecular Biology and Evolution, 35(7), 1588–1598.

[50] Schmeing, T. M. and Ramakrishnan, V. (2009) What recent ribosome structures have revealed about the mechanism of translation. Nature, 461(7268), 1234–1242.

[51] Petrov, A. S., Bernier, C. R., Hsiao, C., Norris, A. M., Kovacs, N. A., Waterbury, C. C., Stepanov, V. G., Harvey, S. C., Fox, G. E., Wartell, R. M., et al. (2014) Evolution of the ribosome at atomic resolution. Proceedings of the National Academy of Sciences, 111(28), 10251–10256.

[52] Holvast, E. J., Celik, M. A., Phillips, M. J., and Wilson, L. A. (2024) Do morphometric data improve phylogenetic reconstruction? A systematic review and assessment. BMC Ecology and Evolution, 24(1), 1–16.

[53] Mitteroecker, P. and Schaefer, K. (2022) Thirty years of geometric morphometrics: Achievements, challenges, and the ongoing quest for biological meaningfulness. American journal of biological anthropology, 178, 181–210.

[54] Bock, L. V., Gabrielli, S., Kolář, M. H., and Grubmüller, H. (2023) Simulation of complex biomolecular systems: the ribosome challenge. Annual Review of Biophysics, 52(1), 361–390.

