## Supplemental material for "Detection of prokaryotic-like ribosome exit tunnels within eukaryotic kingdoms"

The Supplemental Information contains:

- Supplementary methods
- Supplementary Tables S1–S2
- Supplementary Figures S1–S10

### S1 Supplementary methods

#### Automated extraction of ribosome exit tunnel geometry

**Landmark Localization** Three landmarks are used to initiate and refine the results of the MOLE cavity extraction algorithm.

**PTC**: The 3D coordinates of the PTC in each structure are taken to be the midpoint between the ends of the universally conserved LSU rRNA motif (1). The motif itself is located in each structure via a fuzzy search against the rRNA sequences (allowing for Levenshtein distance = 2). The nucleotides constituting the motif belong to the tunnel wall arch making their midpoint a convenient place to initialize the tunnel-search probe.

**Constriction site**: It is taken to be the center of the amino acids cluster in uL4 and uL22 that are positioned less than 15 Å apart.

**Exit Site**: The conserved residues in uL23 and eL39 were identified by selecting the residues closest to the terminal end of centerline coordinates with percentage identity greater than 98% after MSA.

**MOLE probe-based tunnel extraction** We only consider residues within 80 Å of the constriction site. This truncation serves to reduce the computation time and is performed in PyMol.

Following this adjustment, we performed a tunnel search algorithm with **PTC** as the origin. This usually yields 5-10 tunnel branches representative of possible algorithm probe paths.

We further use the **constriction site** and **exit port** positions to pick the tunnel branch aligned with the path of the nascent polypeptide chain, the exit tunnel proper.

To refine the exit direction of the tunnel, a secondary tunnel search is subsequently executed.

**Refinement** Potential inaccuracies in the tunnel geometry near the exit port, such as bending or narrowing, were noted. To address these potential discrepancies, a refinement step was introduced, involving a pairwise comparison of the tunnel identified in the secondary search against all tunnels extracted in the initial search. This comparison leveraged both centerline coordinates and the radii at the exit port of each tunnel. Through this refined process, the tunnel exhibiting the highest degree of similarity in the initial search was selected as the final structure.

To facilitate the comparative analysis of structural variations among tunnels, the geometric representation of each tunnel was simplified to radial variations along the centerline by arc length parameterization using Python. Each tunnel was truncated by 5 Å adjacent to the PTC to remove the region that is sensitive to radius fluctuations. Additionally, a B-spline function with a degree of 7 was applied using Python for smoothing the radial profiles. In the radial plots, the first constriction site was identified by localizing the midpoint of amino acid (aa) residues at aligned position 90 of uL4 protein and position 35 of uL22 for all species using Pymol. For eukaryotes, the second constriction site was located at the midpoint of aa residues at aligned position 104 of uL4 protein and position 21 of uL22. For bacteria and archaea (including the special case of *G.lamblia*), the second constriction site, formed by the structure of rRNA and uL22, was identified as the second trough in the radial plots.

#### Modified sequence similarity assessment

To compare the similarity between the ribosomal protein sequences, we adapted the method proposed by Mervat *et al.* (2), but also accounted for the ambiguous residue "X" in consensus sequences and gaps introduced in MSA. Each type of aa residue $x$ in a protein sequence is represented by intensity $Y_x(i)$ and intensity level $A_x(i)$. The intensity $Y_x(i)$ depends on the abundance and position of each aa:

$$Y_x(i) = f_x i$$

where $f_x$ is the frequency of amino acid $x$ in the sequence, $i$ is the position in an aligned sequence.

The intensity level $A_x(i)$ of each aa is calculated by:

$$A_x(i) = \begin{cases} |10 \ln \dfrac{Y_x(i)}{N}|, & \text{if the element x exits at the position i} \\ 0, & \text{if the element x does not exit at the position i} \end{cases}$$

where $N$ is the number of residues in protein sequence. The combined intensity level of the protein sequence $A_t(i)$ is calculated by the summation of the 22 intensity levels' vectors $A_x(i)$ of the protein sequence:

$$A_t(i) = \sum_{x=1}^{22} A_x(i)$$

The similarity between protein sequences were quantified by evaluating the Euclidean distance among protein sequence descriptor (2), which is a vector composed of the mean $\overline{A}_t$ and standard deviation $SA_t$ of the combined intensity level of the protein sequence introduced by Mervat *et al.* (2):

$$\overline{A}_t = \frac{\sum_{i=1}^{i=N} A_t(i)}{N}$$

$$SA_t = \frac{\sqrt{A_t(i) - \overline{A}_t}^2}{N-1}$$

#### Robustness to the selection of representative tunnel

The resolution of ribosomes ranges from 2 Å to 12 Å, where the choice of relatively low-resolution structures might bias the clustering result. To assess the robustness of our approach in selecting representative tunnels, we used replicate structures from *S.cerevisiae*, which has the most samples and a broad resolution range. Overall, selecting structures close to the cluster centroid did not lead to significant changes in the grouping pattern independent of resolution, demonstrating our tunnel selection strategy is robust to low-resolution data (see Figure S2).

### S2 Supplementary Tables

**Table S1.** Ribosome structures selected with highest resolution for each species

| Domain | Kingdom | Species | Structure selected | Structure with highest resolution |
|---|---|---|---|---|
| Archaea | Methanobacteriati | *H.marismortui* | 4V9F / 2.40 Å (X-ray) | - |
| | | *P.furiosus* | 4V6U / 6.60 Å (EM) | - |
| Bacteria | Bacillati | *B.subtilis* | 8BUU / 2.90 Å (EM) | - |
| | | *E.faecalis* | 7P7T / 2.90 Å (EM) | - |
| | | *L.lactis* | 5MYJ / 5.60 Å (EM) | - |
| | | *M.smegmatis* | 5O60 / 3.18 Å (EM) | - |
| | | *M.tuberculosis* | 7MSZ / 3.10 Å (EM) | 7S0S / 3.05 Å (EM) |
| | | *S.aureus* | 6HMA / 2.65 Å (EM) | 6S0Z / 2.30 Å (EM) |
| | Pseudomonadati | *A.baumannii* | 7RYH / 2.43 Å (EM) | 7RYG / 2.38 Å (EM) |
| | | *E.coli* | 6WD4 / 3.70 Å (EM) | 7K00 / 1.98 Å (EM) |
| | | *F.johnsoniae* | 7JIL / 2.80 Å (EM) | - |
| | | *P.aeruginosa* | 7UNV / 2.70 Å (EM) | 7UNW / 2.60 Å (EM) |
| | Thermotogati | *D.radiodurans* | 5DM6 / 2.90 Å (X-ray) | - |
| | | *T.thermophilus* | 4W29 / 3.80 Å (X-ray) | 4Y4O / 2.30 Å (X-ray) |
| Eukarya | Animalia | *D.melanogaster* | 6XU8 / 3.00 Å (EM) | - |
| | | *D.rerio* | 7OYB / 2.40 Å (EM) | - |
| | | *H.sapiens* | 4UG0 / 3.60 Å (EM) | 7QWQ / 2.83 Å (EM) |
| | | *M.musculus* | 7CPU / 2.82 Å (EM) | - |
| | | *O.cuniculus* | 6P5N / 3.20 Å (EM) | 6SGC / 2.80 Å (EM) |
| | | *R.norvegicus* | 7QGG / 2.86 Å (EM) | - |
| | Fungi | *C.albicans* | 7Q08 / 2.56 Å (EM) | 7PZY / 2.32 Å (EM) |
| | | *E.cuniculi* | 7QEP / 2.70 Å (EM) | - |
| | | *K.lactis* | 4V91 / 3.70 Å (EM) | 6UZ7 / 3.60 Å (EM) |
| | | *S.cerevisiae* | 4U3M / 3.00 Å (X-ray) | 7ZW0 / 2.40 Å (EM) |
| | | *S.pombe* | 8EUI / 3.10 Å (EM) | 8EUG / 2.80 Å (EM) |
| | Plantae | *T.aestivum* | 4V7E / 5.50 Å (EM) | - |
| | Protista | *G.lamblia* | 8BTR / 3.25 Å (EM) | 7PWG / 2.75 Å (EM) |
| | | *L.donovani* | 6AZ3 / 2.50 Å (EM) | - |
| | | *P.falciparum* | 3JBO / 5.80 Å (EM) | 5UMD / 3.20 Å (EM) |
| | | *S.lophii* | 7QCA / 2.79 Å (EM) | - |
| | | *T.cruzi* | 5T5H / 2.54 Å (EM) | - |
| | | *T.gondii* | 5XXB / 3.17 Å (EM) | - |
| | | *T.thermophila* | 7OLC / 2.90 Å (EM) | - |
| | | *T.vaginalis* | 5XY3 / 3.20 Å (EM) | - |

**Table S2.** Taxonomic tree of the species of interest in our study

| Group | Species | Supergroup | Phylum | Family | Genus |
|---|---|---|---|---|---|
| A | *G.lamblia** | Excavata | Fornicata | Hexamitidae | Giardia |
| | *G.muris* | | Fornicata | Hexamitidae | Giardia |
| | *Trepomonas sp.* | | Fornicata | Hexamitidae | Trepomonas |
| | *S.salmonicida* | | Fornicata | Hexamitidae | Spironucleus |
| B | *T.vaginalis** | Excavata | Parabasalia | Trichomonadidae | Trichomonas |
| | *T.foetus* | | Parabasalia | Tritrichomonadidae | Tritrichomonas |
| C | *L.donovani** | Excavata | Euglenozoa | Trypanosomatidae | Leishmania |
| | *T.cruzi** | | Euglenozoa | Trypanosomatidae | Trypanosoma |
| | *T.brucei* | | Euglenozoa | Trypanosomatidae | Trypanosoma |
| | *L.major* | | Euglenozoa | Trypanosomatidae | Leishmania |
| | *S.culicis* | | Euglenozoa | Trypanosomatidae | Strigomonas |
| | *H.muscarum* | | Euglenozoa | Trypanosomatidae | Herpetomonas |
| | *Phytomonas sp.* | | Euglenozoa | Trypanosomatidae | Phytomonas |
| | *T.vivax* | | Euglenozoa | Trypanosomatidae | Trypanosoma |
| | *L.lainsoni* | | Euglenozoa | Trypanosomatidae | Leishmania |
| | *C.expoeki* | | Euglenozoa | Trypanosomatidae | Crithidia |
| | *L.seymouri* | | Euglenozoa | Trypanosomatidae | Leishmania |
| | *L.panamensis* | | Euglenozoa | Trypanosomatidae | Leishmania |
| | *A.deanei* | | Euglenozoa | Trypanosomatidae | Angomonas |
| | *L.pyrrhocoris* | | Euglenozoa | Trypanosomatidae | Leishmania |
| D | *S.lophii** | Amorphea | Microsporidia | Spraguidae | Spraguea |
| | *E.cuniculi** | | Microsporidia | Unikaryonidae | Encephalitozoon |
| | *V.corneae* | | Microsporidia | Nosematidae | Vittaforma |
| | *E.canceri* | | Microsporidia | Enterocytozoonidae | Enterospora |
| | *O.colligata* | | Microsporidia | Ordosporidae | Ordospora |
| | *N.ausubeli* | | Microsporidia | N/A | Nematocida |
| | *N.parisii* | | Microsporidia | N/A | Nematocida |
| | *P.neurophilia* | | Microsporidia | N/A | Pseudoloma |
| E | *T.gondii** | TSAR | Apicomplexa | Sarcocystidae | Toxoplasma |
| | *P.falciparum** | | Apicomplexa | Plasmodiidae | Plasmodium |
| | *T.thermophila** | | Ciliophora | Tetrahymenidae | Tetrahymena |
| | *T.aestivum** | Archaeplastida | Streptophyta | Poaceae | Triticum |
| | *S.pombe** | Amorphea | Ascomycota | Schizosaccharomycetaceae | Schizosaccharomyces |
| | *S.cerevisiae** | | Ascomycota | Saccharomycetaceae | Saccharomyces |
| | *K.lactis** | | Ascomycota | Saccharomycetaceae | Kluyveromyces |
| | *C.albicans** | | Ascomycota | Debaryomycetaceae | Candida |
| | *D.rerio** | | Chordata | Danionidae | Danio |
| | *H.sapiens** | | Chordata | Hominidae | Homo |
| | *O.cuniculus** | | Chordata | Leporidae | Oryctolagus |
| | *M.musculus** | | Chordata | Muridae | Mus |
| | *R.norvegicus** | | Chordata | Muridae | Rattus |
| | *D.melanogaster** | | Arthropoda | Drosophilidae | Drosophila |

Species* with available ribosome structures.

#### S3 Supplementary Figures

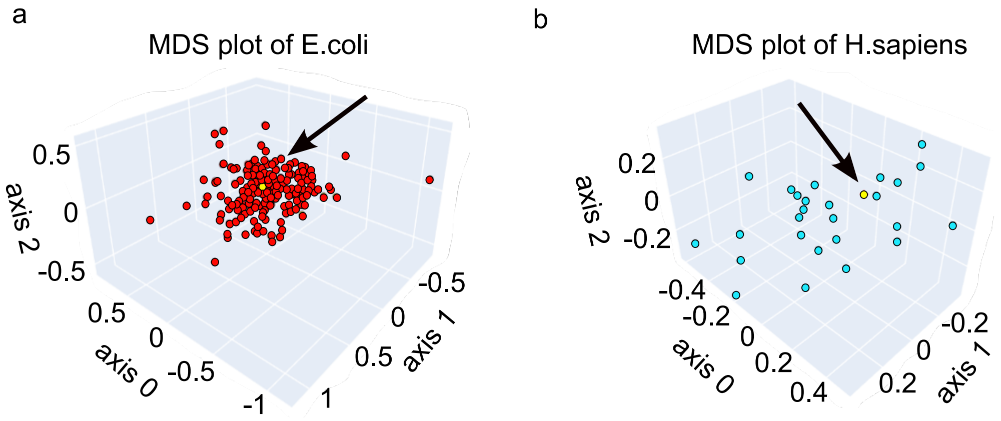


**Figure S1.** Selection of representative structure of **(a)** *E.coli* and **(b)** *H.sapiens*. Each data point represents a single tunnel. The yellow point indicates the one closet to the cluster centroid.

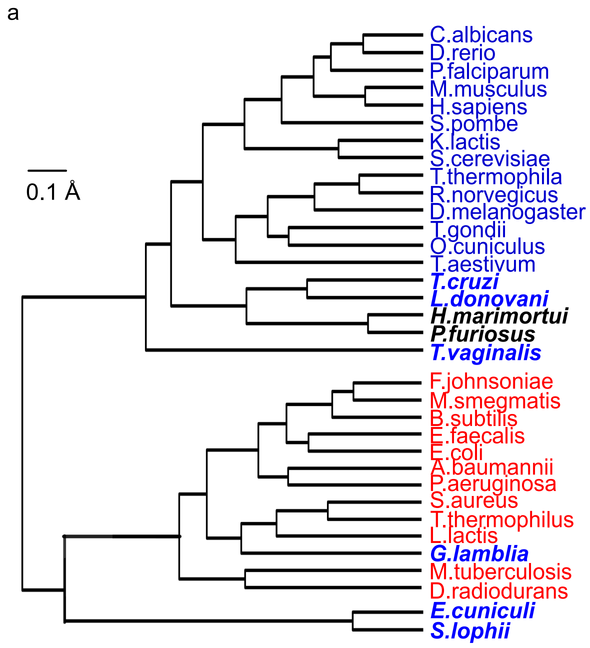


**Figure S2.** Comparison of the hierarchical clustering constructed using: **(a)** the structure closest to the cluster centroid, and **(b)** the highest-resolution structure available for each species.

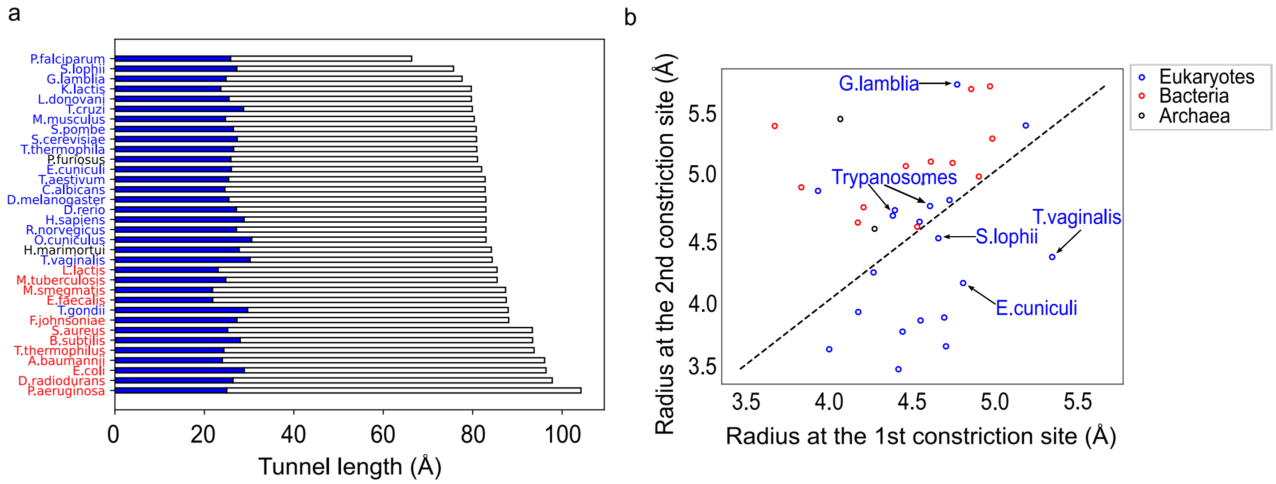


**Figure S3.** Geometric comparison of the ribosome exit tunnel. **(a)** The horizontal bar plot shows the exit tunnel length. The length is decomposed into two subparts, separated by the universally shared constriction site. Bacteria, archaea and eukaryotes are colored by red, black and blue. **(b)** Comparisons of the radii at the two constriction sites. In bacteria, the 'second constriction site' is defined as the second trough in the radial plot. We observe that all bacteria (red dots), archaea (black dots) and some of eukaryotes (blue dots) including *G.lamblia*, *L.donovani* and *T.cruzi* exhibit wider second constriction site. In contrast, the other outlier species *T.vaginalis*, *S.lophii* and *E.cuniculi* have narrower ones.

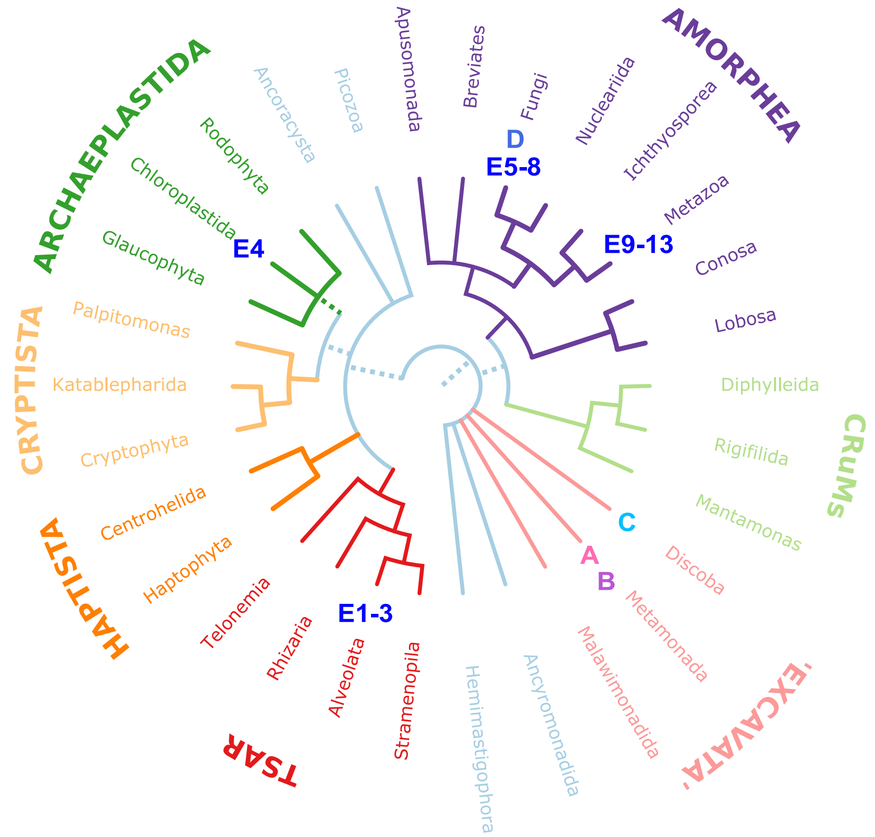


**Figure S4.** Schematic tree of eukaryotes showing the phylogenetic positions of species studied. The colored groupings correspond to the 'supergroups'. This work is inspired by Burki *et al.* (3) and adapted from 'Eukaryote Tree of Life 2020' by Tricholome, licensed under CC-BY-SA 4.0.

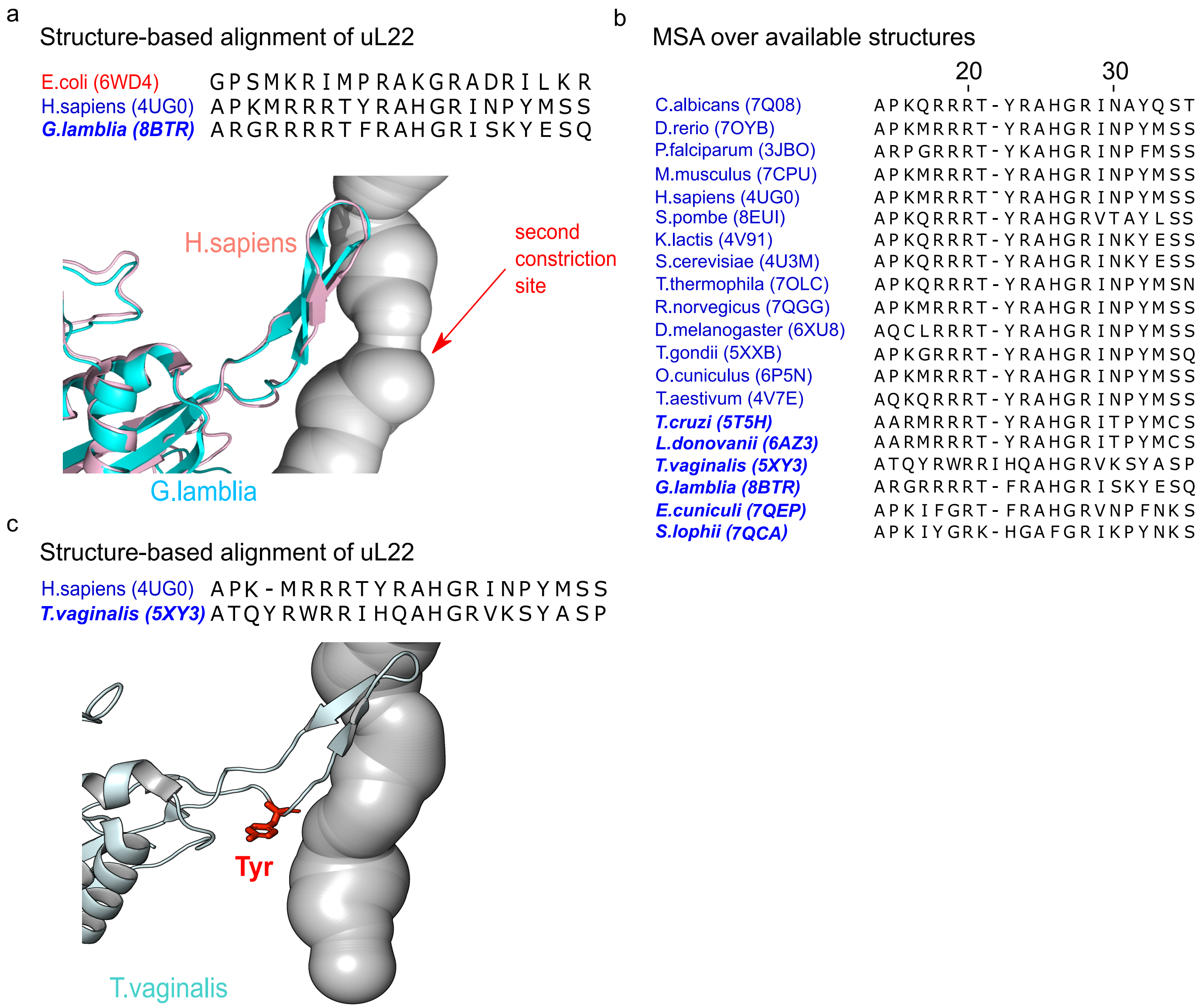


**Figure S5.** The structure of ribosomal protein uL22 has no effect on the constriction sites. **(a)** The structure-based alignment of uL22 from *E.coli*, *H.sapiens*, and *G.lamblia*. The lower panel shows the tunnel structure of *H.sapiens* and the structural superposition of uL22 proteins of *H.sapiens*(pink) an *G.lamblia* (skyblue). **(b)** The sequence alignment of uL22 proteins from all eukaryotic species involved in our hierarchical clustering. Only the amino acids within 15 Å of the tunnel centerline are displayed (from aligned position 14 to 35). Notably, for *T.vaginalis*, I22 is not aligned with other sequences. **(c)** Comparison of the ul22 structures of *H.sapiens* and *T.vaginalis* reveals a Tyr residue in *T.vaginalis* that is not conserved across other eukaryotic species. This Tyr residue protrudes away from the exit tunnel, confirming that this change has no effect on the tunnel structure.

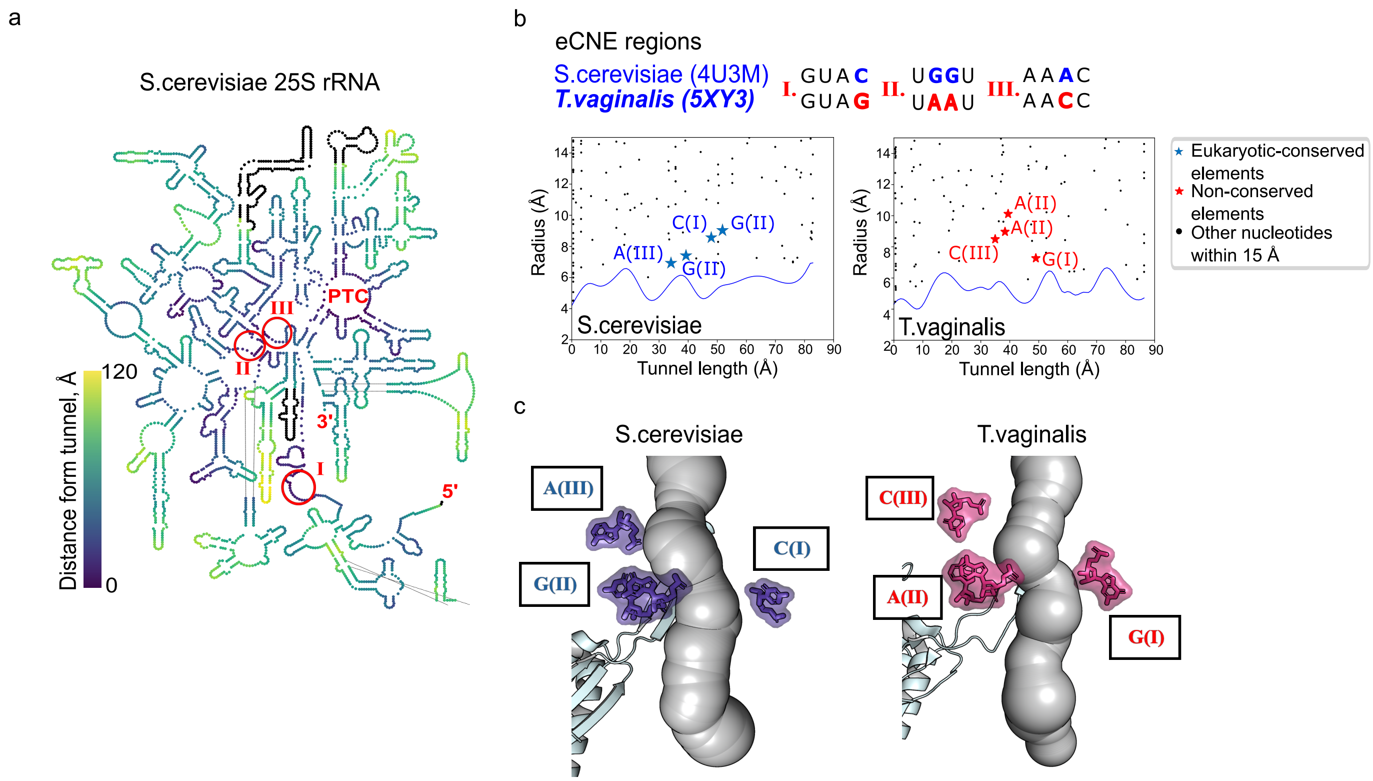


**Figure S6.** Conservation study of LSU rRNA in *T.vaginalis.* **(a)** A map of the secondary structure of the 25S rRNA in *S.cerevisiae*, which served as a structural filter in previous research (1), colored by distance from the tunnel. The map is retrieved from RiboVision database and edited using the module "Color by Data". Three eukaryotic conserved nucleotide elements (eCNEs) regions close to the tunnel within 15 Å are highlighted. **(b)** Alignments of LSU rRNA subsequences from *S.cerevisiae* and *T.vaginalis* in three eCNEs regions. The conserved nucleotides (blue) are mapped into the radial plot of *S.cerevisiae* as blue stars. Non-conserved nucleotides (red) are mapped into the radial plot of *T.vaginalis* as red stars. All other nucleotides within 15 Å of the tunnel centerline are depicted as black circles. Alphabets represent nucleotide elements and roman letters represent region numbers. Regions are numbered in the order of the nucleotide sequence. **(c)** Structures of those rRNA nucleotides in *S.cerevisiae* and *T.vaginalis* and their relative positions to the exit tunnel. Non-conserved nucleotides in *T.vaginalis* are shown in red. We observed notable voids between the mutated nucleotides and the tunnel surface in *T.vaginalis*, indicating that its tunnel structure is shaped by ribosomal proteins uL4 and eL39, rather than rRNAs.

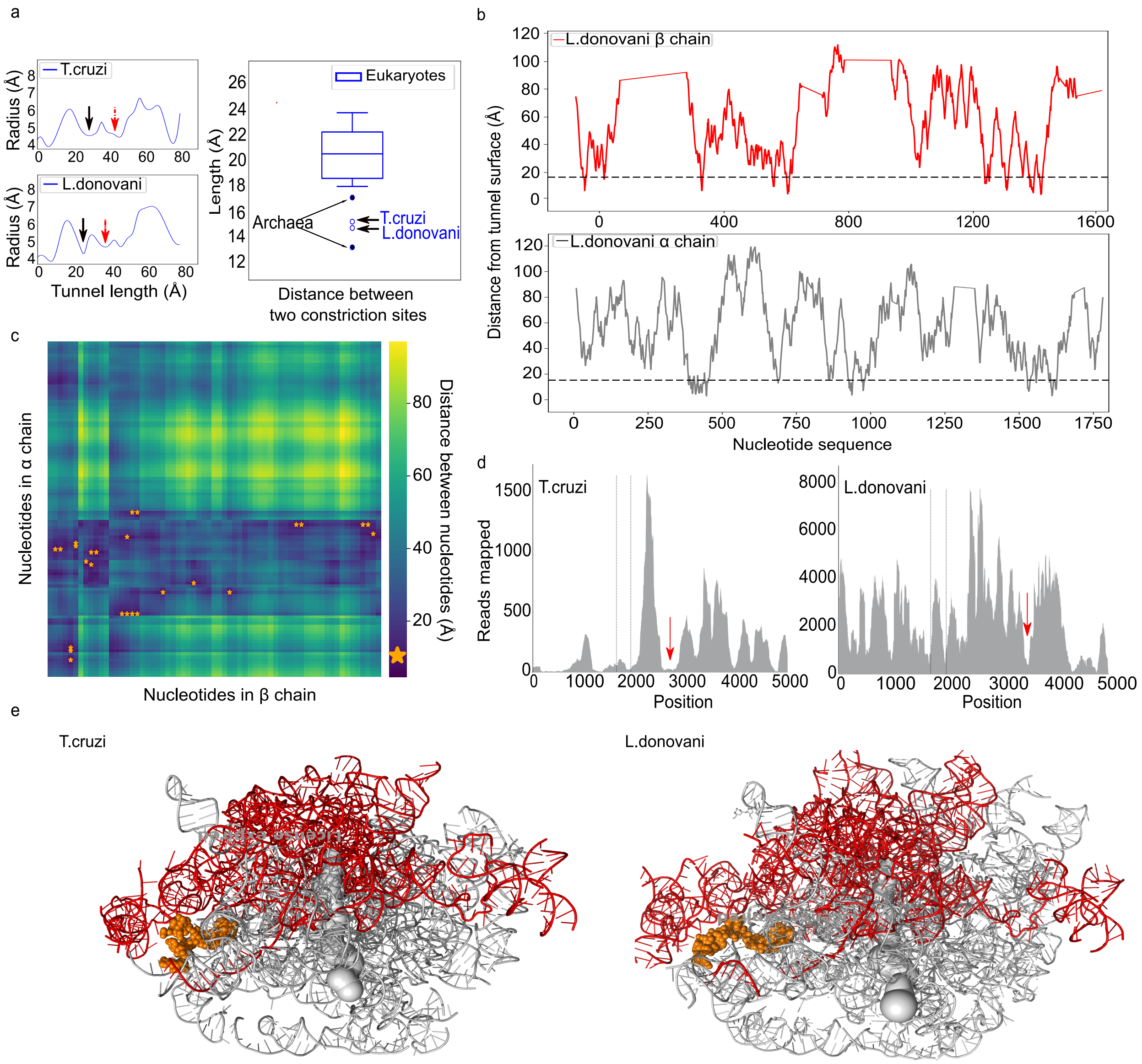


**Figure S7.** The non-homologous break in trypanosomes reduces the distance between constriction sites. **(a)** The left inset displays the radial plots of *T.cruzi* and *L.donovani*, with the first and second constriction sites marked by black and red arrows, respectively. The distances between them as shown in the box plot on the right. **(b)** The distances between the nucleotides in $\beta$ (upper) and $\alpha$ (lower) chains and the tunnel surface for *L.donovani*. The dash lines represent the cutoff distance at 15 Å. **(c)** Heat map of selected nucleotides for *L.donovani*. Nucleotide pairs with distance less than 10 Å are highlighted with orange stars. **(d)** The mapped rRNA readings of *T.cruzi* and *L.donovani* generated by the hidden break detection pipeline. The dash lines represent the positions of two highly conserved 20-mers flanking the hidden break region characterized in protostomes. The red arrows show the break positions in trypanosomes. **(e)** The LSU structures of *T.cruzi* and *L.donovani* with conserved motifs highlighted in orange.

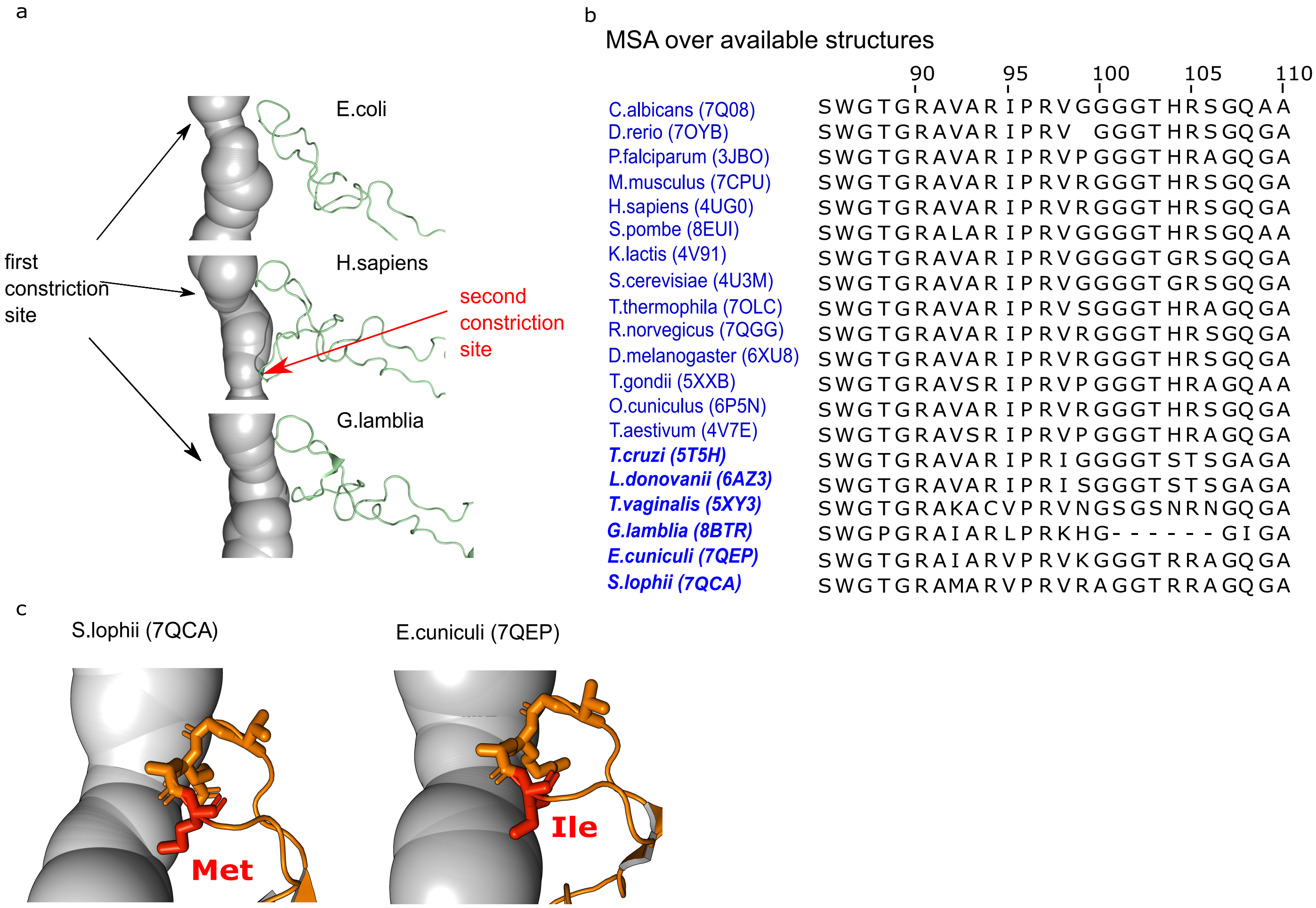


**Figure S8.** A missing segment in uL4 protein sequence of *G.lamblia* induces a wider second constriction site. **(a)** Comparison of the uL4 structure between *E.coli*, *H.sapiens* and *G.lamblia*. **(b)** Sequence alignment of uL4 proteins from all eukaryotic species involved in our hierarchical clustering. Only the amino acids close to the two constriction sites of the tunnel within 15 Å are displayed (from aligned position 85 to 110). **(c)** At aligned position 92, *S.pombe*, *G.lamblia*, and *E.cuniculi* have non-conserved amino acids L (Leu) and I (Ile). These non-conserved amino acids have no effect on the constriction site due to the short length of their side chain. In contrast, in *E.cuniculi*, the non-conserved amino acid is M (Met), which induce additional distortions between the two constriction sites. This tunnel geometric feature can be observed in its radial plot.

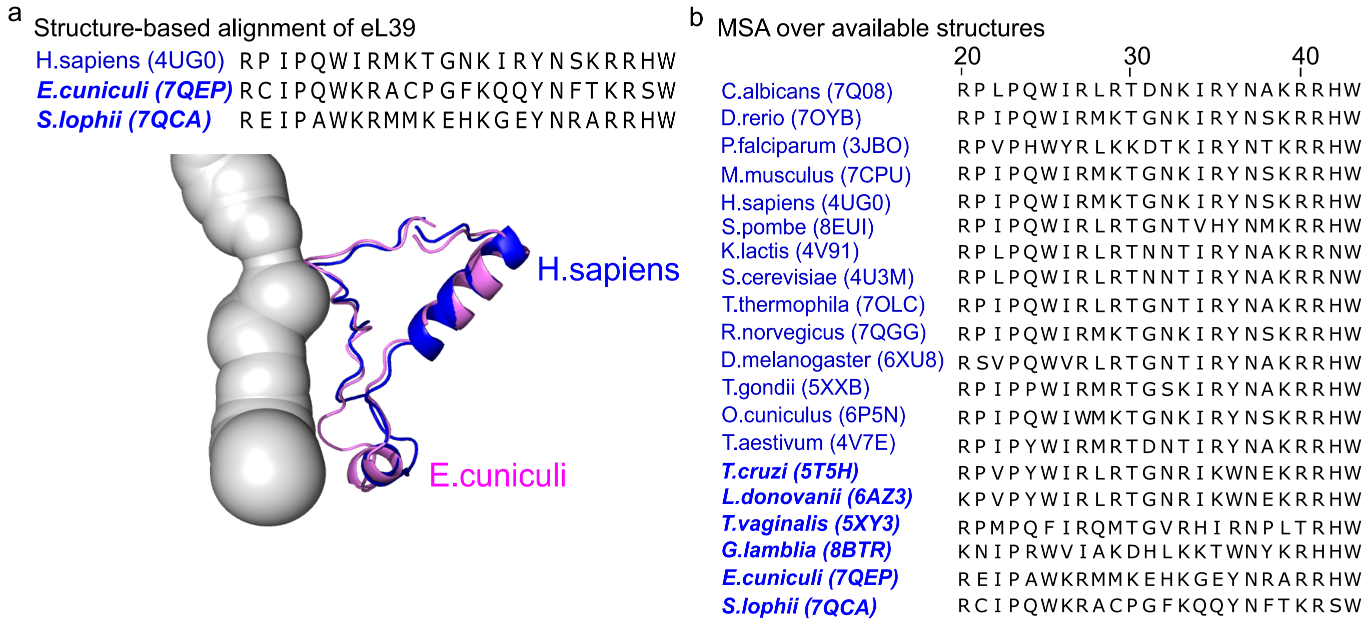


**Figure S9.** The structure of ribosomal protein eL39 has no effect on geometric variation in the lower region of eukaryotic tunnels. **(a)** Structure-based alignment of eL39 from *H.sapiens* and two outliers *E.cuniculi* and *S.lophii*. The lower panel shows the tunnel structure of *H.sapiens* and the structural superposition of eL39 proteins of *H.sapiens* (blue) an *E.cuniculi* (magenta). **(b)** Sequence alignment of eL39 proteins from all eukaryotic species involved in our hierarchical clustering. Only the amino acids within 15 Å of the tunnel centerline are displayed (from aligned position 20 to 43).

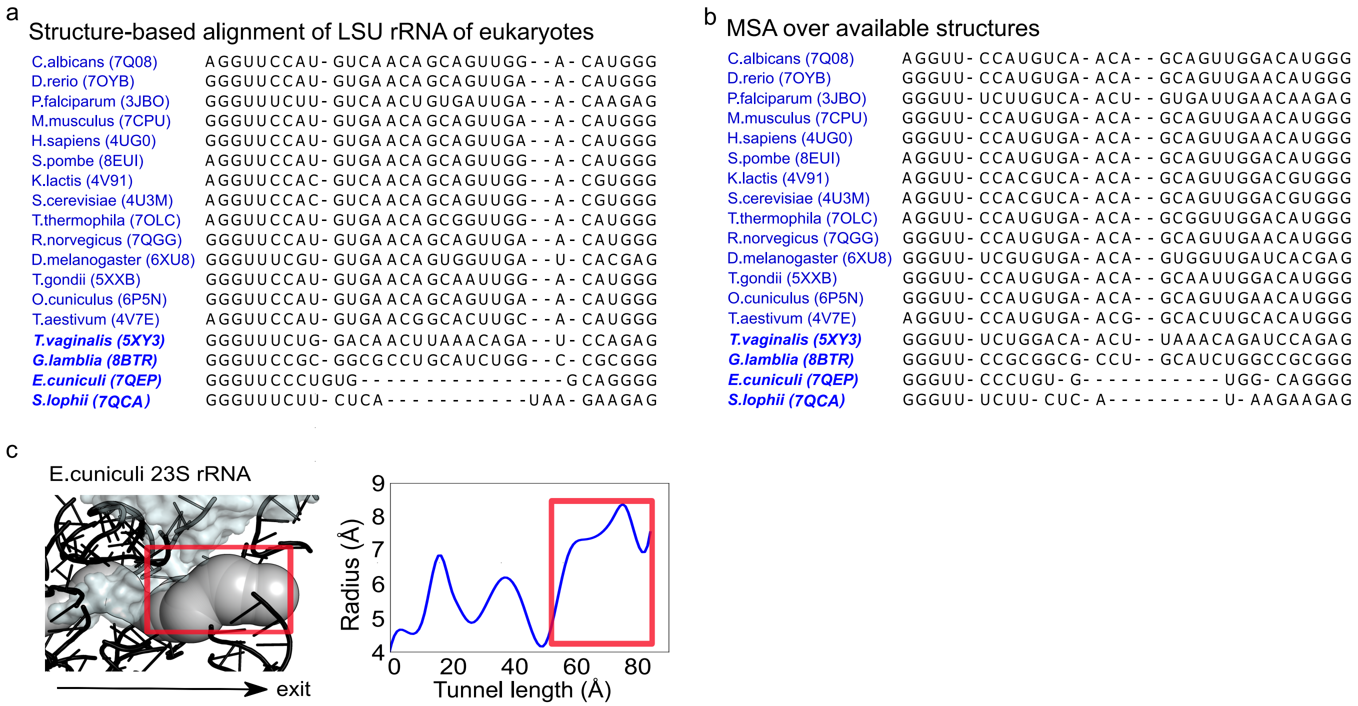


**Figure S10.** **(a)** Structure-based alignment and **(b)** multiple sequence alignment of LSU rRNA from all eukaryotic species involved in our hierarchical clustering except for two trypanosomes. The subsequences near the tunnel exit port are truncated and only *E.cuniculi* and *S.lophii* are observed with missing segments in two types of alignments. **(c)** The LSU rRNA structure and associated radial plot of *E.cuniculi*. The missing rRNA part is indicated by red box.

### REFERENCES


1. Doris, S. M., Smith, D. R., Beamesderfer, J. N., Raphael, B. J., Nathanson, J. A., and Gerbi, S. A. (2015) Universal and domain-specific sequences in 23S–28S ribosomal RNA identified by computational phylogenetics. Rna, **21**(10), 1719–1730.

2. Abo-Elkhier, M. M., Abd Elwahaab, M. A., Abo El Maaty, M. I., et al. (2019) Measuring similarity among protein sequences using a new descriptor. BioMed research international, **2019**.

3. Burki, F., Roger, A. J., Brown, M. W., and Simpson, A. G. (2020) The new tree of eukaryotes. Trends in ecology & evolution, **35**(1), 43–55.
